# Comprehensive genomic analysis reveals dynamic evolution of endogenous retroviruses that code for retroviral-like protein domains

**DOI:** 10.1101/628875

**Authors:** Mahoko Takahashi Ueda, Kirill Kryukov, Satomi Mitsuhashi, Hiroaki Mitsuhashi, Tadashi Imanishi, So Nakagawa

**Affiliations:** Department of Molecular Life Science, Tokai University School of Medicine, Isehara, Kanagawa 259-1193, Japan; Micro/Nano Technology Center, Tokai University, Hiratsuka, Kanagawa 259-1292, Japan; Department of Human Genetics, Yokohama City University Graduate School of Medicine, Yokohama, Kanagawa 236-0004, Japan; Department of Applied Biochemistry, School of Engineering, Tokai University, Hiratsuka, Kanagawa 259-1292, Japan; Institute of Medical Sciences, Tokai University, Isehara, Kanagawa 259-1193, Japan

**Keywords:** endogenous retrovirus, transposon exaptation, co-option, de novo gene

## Abstract

Endogenous retroviruses (ERVs) are remnants of ancient retroviral infections of mammalian germline cells. A large proportion of ERVs lose their open reading frames (ORFs), while others retain them and become exapted by the host species. However, it remains unclear what proportion of ERVs possess ORFs (ERV-ORFs), become transcribed, and serve as candidates for co-opted genes. Hence, we investigated characteristics of 176,401 ERV-ORFs containing retroviral-like protein domains (*gag, pro, pol*, and *env*) in 19 mammalian genomes. The fractions of ERVs possessing ORFs were overall small (∼0.15%) although they varied depending on domain types as well as species. The observed divergence of ERV-ORF from their consensus sequences suggested that a large proportion of ERV-ORFs either recently or anciently inserted themselves into mammalian genomes. Alternatively, very few ERVs lacking ORFs were found to exhibit similar divergence patterns. To identify ERV-ORFs transcribed as proteins, we compared ERV-ORFs with various multi-omics data including transcriptome data, trimethylation at histone H3 lysine 36, and transcription initiation sites from 2,834 cell types, and found 408 and 752 ERV-ORFs, accounting for 2-3% of all ERV-ORFs, with high transcriptional potential in humans and mice, respectively. Moreover, many of these ERV-ORFs with transcriptional potential were lineage-specific sequences exhibiting tissue-specific expression. These results suggest a possibility for the expression of uncharacterized functional genes containing ERV-ORFs hidden within mammalian genomes. Together, our analyses suggest that more ERV-ORFs may be co-opted in a host-species specific manner than we currently know, which are likely to have contributed to mammalian evolution and diversification.

## Introduction

Transposable elements (TEs), also known as “jumping genes”, constitute large portions of mammalian genomes. It has been reported that up to 70%, of the human genome is originated from TEs (International Human Genome Sequencing Consortium 2001; de Koning 2011; Smit et al. 2013). Generally, TEs are categorized as junk DNA (Smit 1999); however, many studies have shown that TEs have contributed greatly to mammalian evolution. Furthermore, TEs promote rearrangements of chromosomal DNA (McVean 2010), and can become sources of both coding and regulatory sequences during the evolution of host genomes (Thornburg et al. 2006; Nishihara et al. 2016; Chuong et al. 2016; Diwash et al. 2017). In particular, specific types of long terminal repeat (LTR) retrotransposons, including endogenous retroviruses (ERVs), have been shown to develop the ability to function as genes in several mammalian tissues (Ono et al. 2006; Matsui et al. 2011; Nakaya et al. 2013; Pastuzyn et al. 2018; Ashley et al. 2018). One of the most well-studied ERV-co-opted genes is syncytin. Specifically, the human syncytin-1 (*ERVW-1*) gene, which is derived from an envelope (*env*) gene in human endogenous retrovirus type W (*HERV-W*), was exapted for cell fusion during placental development (Mi et al. 2000; Dupressoir et al. 2012; Lavialle et al. 2013; Bolze et al. 2017). Interestingly, many of the molecular functions of ERV-co-opted genes remain the same as those in viruses (Mi et al. 2000; Pastuzyn et al. 2018; Ashley et al. 2018).

ERVs are thought to be derived from retroviruses established in the germ line of various organisms in the past. In the human genome, ERVs occupy approximately 8% of the human genome (International Human Genome Sequencing Consortium 2001; de Koning 2011; Smit et al. 2013). Many ERVs lose their open reading frames (ORFs) by accumulating deletions or mutations after integration. Thus, it is unclear what percentages of ERVs in mammalian genomes possess retroviral-like protein ORFs and have maintained high transcriptional potential. Thus far, genome-wide comparative analyses using epigenomic and transcriptomic data have examined the regulatory regions of ERVs (Ito et al. 2017); however, they have not examined characterized ERV-derived protein-coding sequences. Recently, many whole transcriptome sequencing (RNA-Seq) datasets have been accumulated in public databases. One of the most comprehensive RNA-Seq datasets was generated by the genotype-tissue expression (GTEx) study using 31 human tissues (Carithers et al. 2015). The data was used to assemble a similar pipeline in the Comprehensive Human Expressed SequenceS (CHESS) project and served to generate 20,352 potential protein-coding genes, and 116,156 novel transcripts in the human genome (Pertea et al. 2018). A comparative analysis study focusing on the protein-coding region of ERVs using the new transcript data, in combination with cap analysis of gene expression (CAGE) data (Lizio et al. 2015), and epigenomic data will provide new insight into the transcriptional potential, and functionality of these regions.

To understand the characteristics of possible protein-coding ERVs that possess ORFs for retroviral-like protein domains (ERV-ORFs) in mammalian genomes, we comprehensively examined ERV-ORFs in 19 mammalian species. We previously developed a database called gEVE (http://geve.med.u-tokai.ac.jp) containing 176,401 ERV-ORFs containing at least one functional viral gene motif found in 19 mammalian genomes (Nakagawa and Takahashi 2016). In this study, we analyzed ERV-ORFs in the gEVE database using various transcriptome and epigenomic data in humans and mice including the abovementioned data. Several systematic searches were performed to obtain ERV-ORFs at the domain level; these studies were designed to detect ERVs in the human genome (Paces et al. 2002; de Parseval et al. 2003; Villesen et al. 2004; Tokuyama et al. 2018). Most of these studies were limited to protein-coding ERVs in humans or mice, and primarily examined ERV sequences that contained nearly full-length ORFs or specific domains. Importantly, ERV-derived genes have also been described that do not possess intact ORFs yet continue to play important roles in specific situations (Sugimoto et al. 2013). Moreover, multi-exon genes have been identified that contain exons partially derived from ERV sequences (Sela et al. 2007). Note that ERV-ORFs stored in the gEVE database do not always possess ORFs starting with an initiation codon (i.e. an ATG triplet); however, all must contain viral-like protein domains that were predicted using Hidden Markov models (HMMs). The identified ORFs were primarily from four ERV genes [*gag* (viral core proteins), *pro* (proteases), *pol* (polymerase), and *env* (envelope)], all of which are commonly found in viral amino acid sequences. In this study, we performed genome-wide comparisons on the number, divergence, genomic distribution, and transcriptional potential for each protein domain in the ERV-ORFs of mammals. Our findings are expected to improve the understanding of the evolution and potential roles of unannotated ERV-derived genes.

## Results

### Characteristics of ERV-ORFs in 19 mammalian genomes

To characterize ERVs that are expressed as proteins in mammalian species, we first compared 176,401 possible protein-coding ERV-ORFs from 19 mammalian species genomes obtained from the gEVE database (Nakagawa and Takahashi 2016). ERV-ORFs include ORFs of ≥80 amino acid (aa) residues that encode domains in retroviral genes such as *gag, pro, pol*, and *env* genes. The number of ERV-ORFs vary among mammalian genomes (Figure 1A), as we have reported previously (Nakagawa and Takahashi 2016). In each genome, the percentage of possible protein-coding ERVs relative to the total ERVs, were found to range from 0.05% to 0.15% (Figure 1A). Specifically, within mice and cows, the proportion of detectable ERV-ORFs was higher compared to other species. Moreover, the proportion of ERV-ORFs was significantly correlated with the total ERV genome length (ERV-ORF: *r* = 0.71, *p* < 5.51 × 10^−4^). Distributions of the ERV-ORF lengths among mammals are shown in Figures S1A-S1C.

**Figure 1.**
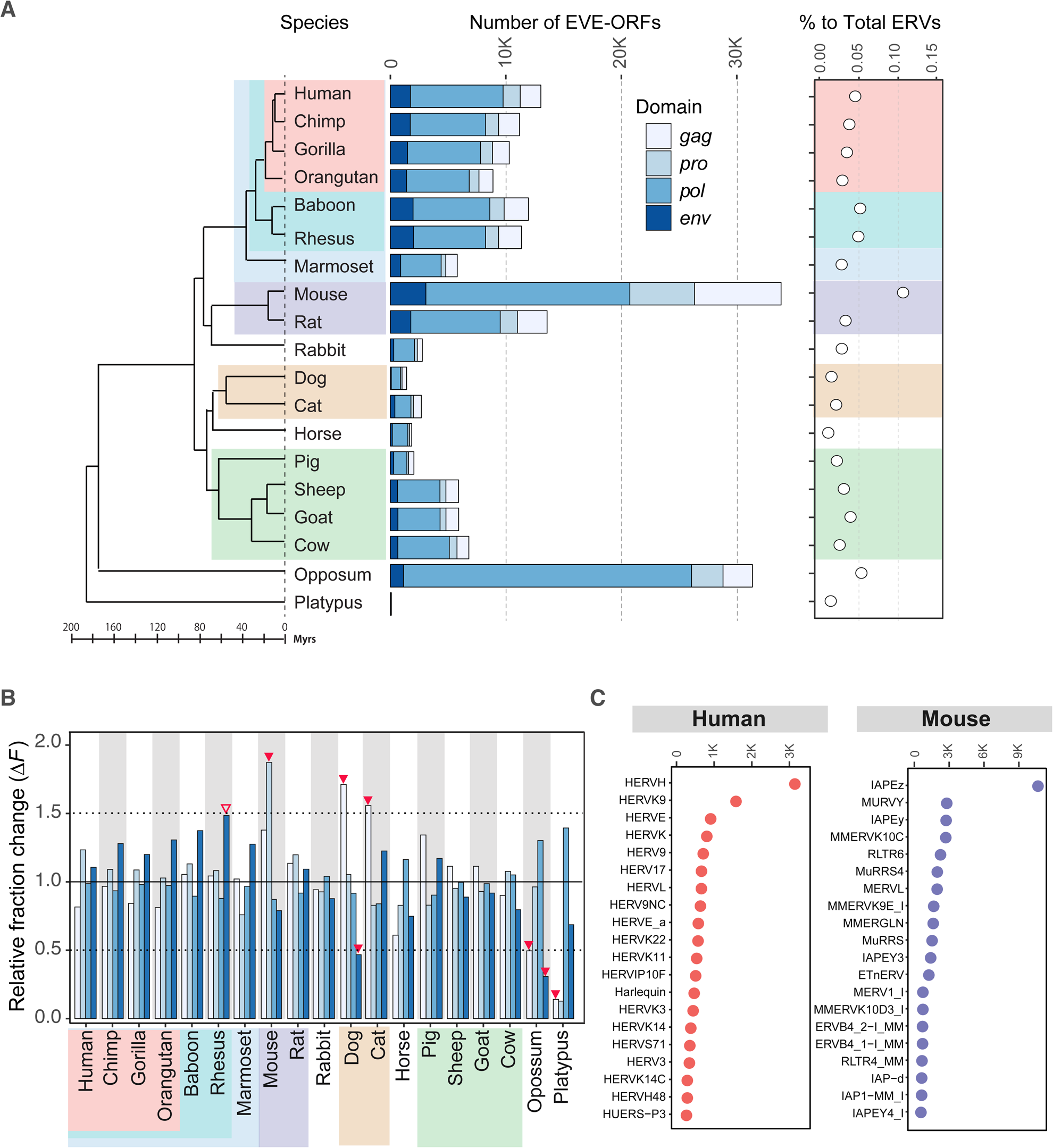
Number of identified ERVs and Repbase annotations in 19 mammalian species. A) The mammalian phylogeny showing the number of identified ERV-ORFs. Background colors for each species’ name indicate that they have shared species classification equal or below the level of order. Bar colors represent each domain. The proportion of ERV-ORFs identified compared to all ERV sequences in each species, are shown on the right. Chimp, chimpanzee; Rhesus, rhesus macaque monkey. B) Relative change in the number of each protein domain identified compared to the mammalian average (Δ*F*) is shown on the y-axis. Colors of protein domains and mammalian classification are the same as those used in panel A. Any Δ*F* that was determined to be more than 1.5 or less than −1.5 are highlighted by red triangles. The Δ*F* for the *env* gene of rhesus macaques, which was approximately 1.5 (1.49), is also highlighted by an open triangle. C) Top 20 Repbase annotations for ERVs in the human and mouse genomes. The x-axis represents the number of ERV-ORFs in each annotated group.

We further compared mammalian ERV-ORFs at the level of gene domains. To reduce the influence of genome qualities on the number of domains detected, we calculated the relative fraction change (Δ*F*) from the mammalian average for each domain fraction (Figure 1B). The most species-specific characteristics observed in the Δ*F* values were determined to be in the *gag* gene, which exhibited a ΔF of more than ±1.5 in several species (Figure 1B). In addition, the *pro* gene, in mice expressed a dramatic increase (Δ*F* = 1.8). Furthermore, platypuses showed quite unique compositions in their *gag* and *pro* genes with very small *ΔF* values (0.14 and 0.12, respectively). Alternatively, the *ΔF* values for *pol* were similar among mammalian species, all of which had expression levels that were determined to be within ±1.5 of each other. Interestingly, although no species showed Δ*F* values greater than ±1.5 in the *env* gene, the values were relatively higher within non-human primates, specifically within rhesus macaque, which had a Δ*F* of nearly 1.5.

We next examined which ERV groups from the Repbase database (Bao et al. 2015) were enriched in human and mice ERV-ORFs (herein we employ the term “group” rather than the Repbase classification of “families/sub-families” for ERVs to avoid confusion with species classification). A large proportion of the detected ERVs in humans were found to be HERV-H and HERV-K, and IAPE (Intracisternal A-type Particles elements with an Envelope) in mice (Figure 1C). HERV-H represents the ancient HERV group, which was inserted into the common ancestral genome of simians and prosimians (Bénit et al. 2003), while HERV-K is described as containing younger HERVs (Bannert and Kurth, 2006; Subramanian et al. 2011). In addition, the IAPE in mice is one of the ERV groups that continue to be active in mice (Mouse Genome Sequencing Consortium, 2002).

We identified divergent fragments of ERV-ORFs using the Repbase consensus sequences, and compared them with ERVs excluding ERV-ORF regions (hereafter called non-ERV-ORFs). We hypothesized that newer elements would be closer to the consensus, and older ones would be further. Thus, we were able to estimate the age of ERV-ORFs by comparing the divergence spectra to their consensus sequences. We first determined modes of ERV divergence based on kernel density estimates (KDE). The modes of divergence were quite different from those of non-ERV-ORFs which were found to be approximately 20% in all mammalian genomes (Figure 2). The ERV-ORFs showed clear bimodal distributions in many mammalian species, with modes of approximately 2 - 7% (younger) and 30 - 33% (older). It should be noted that in specific species the distribution patterns revealed weak or biased bimodality. Specifically, the bimodality was found to be weak and biased in dogs, sheep, and opossum, and small divergences were observed in humans. Although these distribution patterns were relatively similar among closely related species, such as non-human primates and rodents, overall, the distributions varied significantly between species (Figure 2).

**Figure 2.**
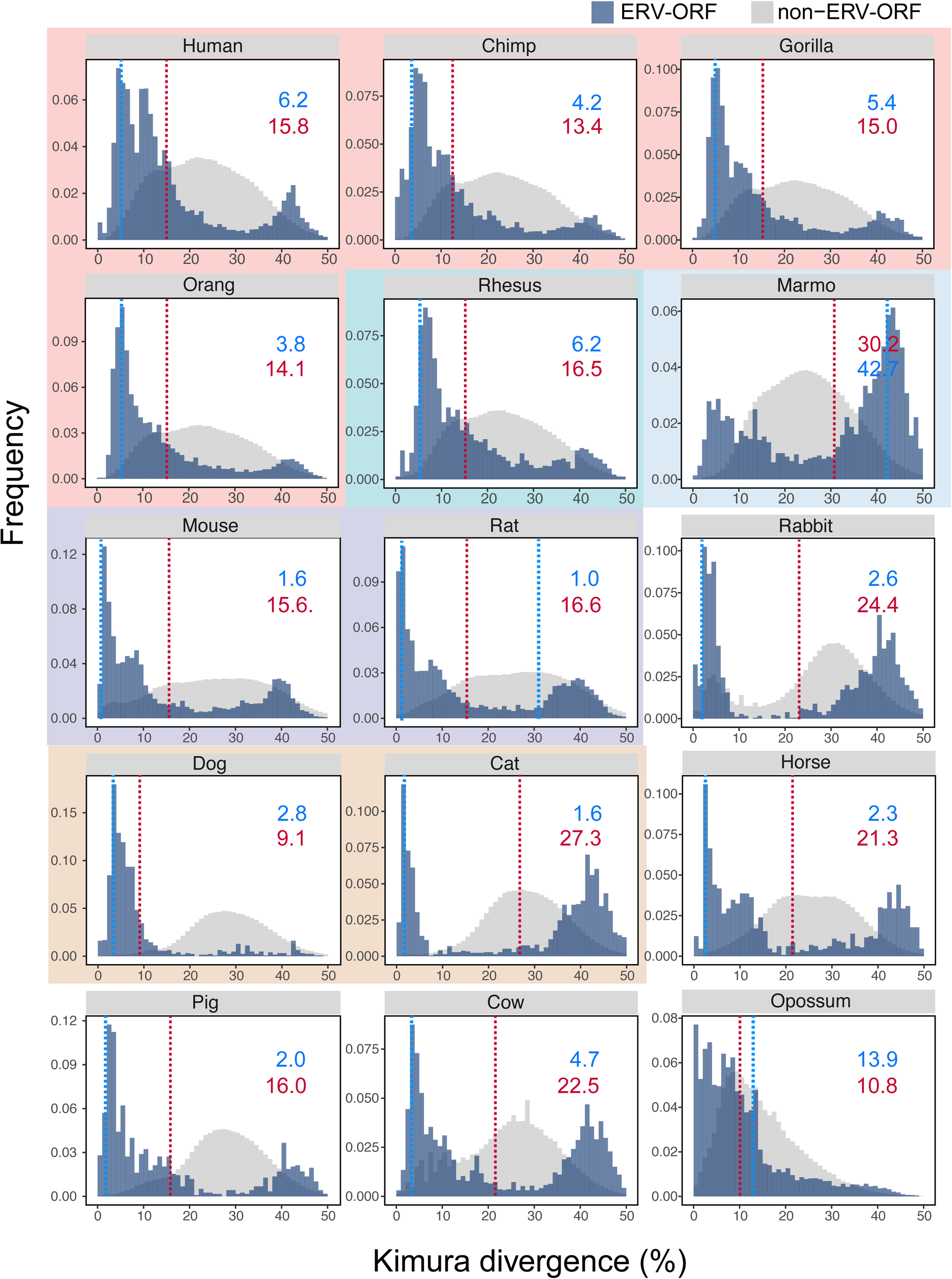
Divergence from Repbase consensus sequences. Divergence frequencies for the consensus sequences in ERV-ORF (blue) and non-ERV-ORF (gray). Background colors for each panel represent the corresponding mammalian classification as indicated in Figure 1A. For each species, the mode and average values for ERV divergence are shown in blue and red, respectively.

Finally, we determined how many ERV-ORFs act as raw materials for the assembly of multi-exon genes. Comparison of our observed ERV-ORFs with those in Ensembl gene and protein annotations revealed that 13, and 24 ERV-ORFs were constituents of ordinary multi-exon genes in humans and mice, respectively (Figure S2 and Table S1). It is also intriguing that large numbers of these multi-exon genes were found in horses (180 genes), sheep (337 genes), and opossum (737 genes). Specifically, within horses and sheep, the number of multi-exon genes containing ERV-ORFs, as predicted by HMM not RepeatMasker (Smit et al. 2013), were quite large (25: horses, 18: sheep, Figure S2).

### Omics analyses of ERV-ORFs

To predict whether ERV-ORFs are expressed as proteins, we performed comparative analyses using a wide-range of transcriptome and histone mark data generated by next-generation sequencing. We first compared observed ERV-ORFs from our study with quantified transcriptome data derived from 31 human tissues generated in the GTEx study (Carithers et al. 2015) from the CHESS database (Pertea et al. 2018). We found that a total of 279 ERV-ORFs overlapped with transcripts in the CHESS database. Of these, 7 ERV-ORFs were found to correspond to genes containing exons derived from ERV sequences predicted only by HMM, not by RepeatMasker (Table S1). The proportion of ERV-ORFs overlapping with CHESS transcripts in each domain were 3.0, 2.9, 3.5, and 4.7 in *gag, pro, pol*, and *env*, respectively. This suggests that the *env* segment was relatively large compared to the other domains, however, this result was not statistically significant (chi-square test *p* < 0.16).

Since ERVs are exclusively expressed during embryogenesis and cell differentiation in mammals (Lin et al. 1999; Grow et al. 2015), the RNA-Seq data acquired by the GTEx project is insufficient to obtain accurate information on ERV-ORFs as it exclusively reflects samples obtained from fully differentiated human tissues. We, therefore, also analyzed RNA-Seq data from human and mouse differentiating myoblasts. The relevant expression data from human primary myoblasts (12 runs) and mouse C2C12 cells (16 runs) were collected from the sequence read archive (SRA) database (see Supplementary Table 2 for details). Following gene mapping and quantification, the expression data of ERV-ORFs having a minimum of ten normalized read counts in the human and mouse myoblasts were extracted. The RNA-Seq data were analyzed from four (Days 0-3) and three (Days 0, 3, 6) time points after differentiation had begun in humans and mice, respectively. Principal component analysis (PCA) plots successfully captured the difference in ERV-ORF expression at each time point (Figures 4A and S3). From these plots, 51 and 586 ERV-ORFs were identified as having a minimum of ten normalized read counts in all samples at the same differentiation stage in humans and mice, respectively. In both species, a number of ERV-ORFs exhibited high expression levels throughout the entire differentiation process, including ERV3-1 in humans, and AH000833 and AI506816 in mice. (Figure 3B). Other known genes/transcripts such as Peg10, AC016577 and PPHLN1 in humans, and syncytin-A and Asprv in mice were also detected, however, their expressions were low. Many ERV-ORFs detected in the myoblast differentiation were unreported transcripts, derived from all four ERV domains of ERV1, ERVK, and ERVL, some of which exhibited stage-specific expression patterns (Figure 3B and Figure S5). It is noteworthy to mention that in ERV-ORFs expressed during human muscle cell differentiation, only eight were also identified in GTEx transcripts.

**Figure 3.**
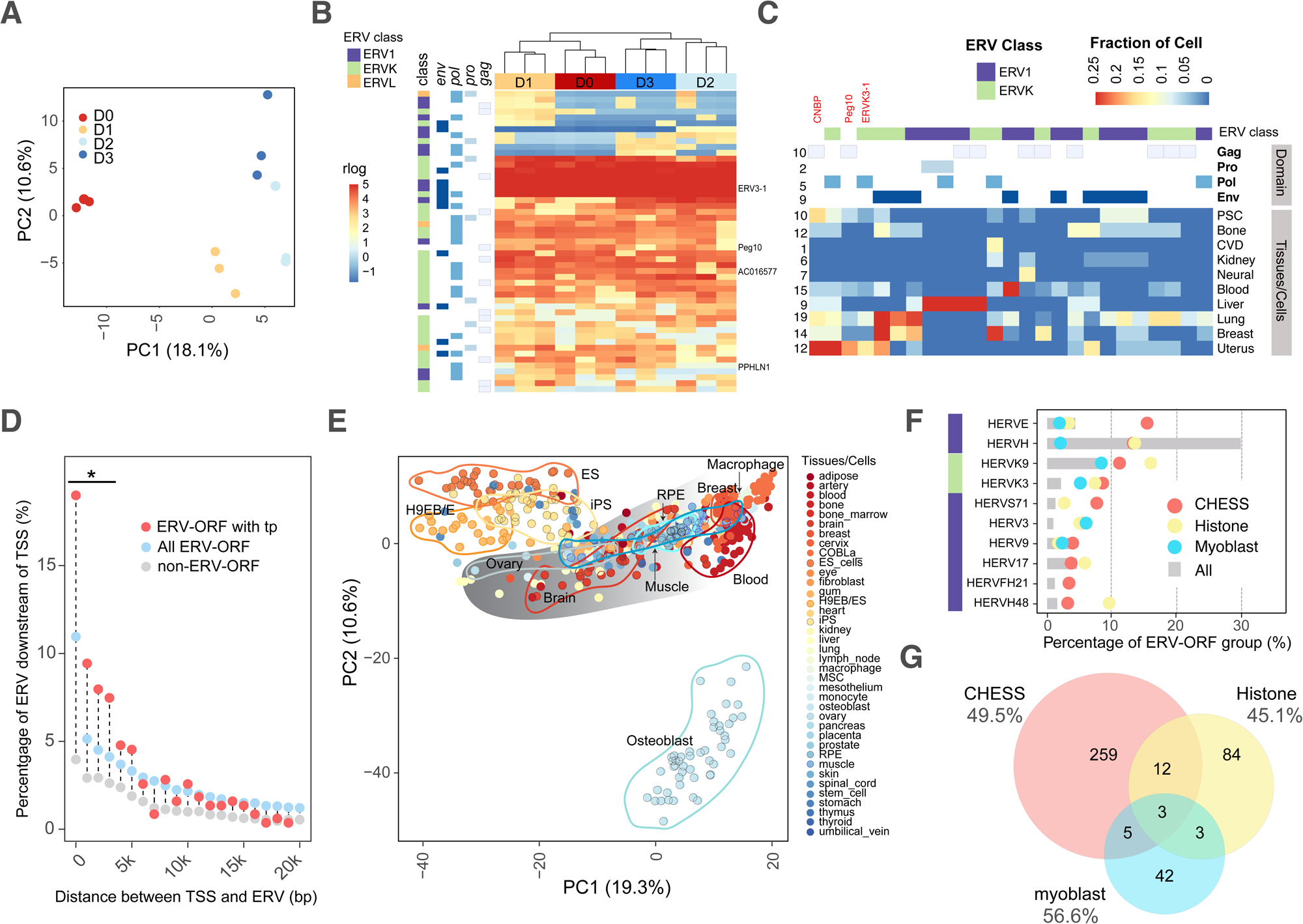
Overlapping human ERV-ORFs with functional data in various tissues and cell lines. (A) PCA plot illustrating the ERV-ORFs expressed during the four stages of human primary myoblast differentiation (three replicates in each stage) for the first two principal components (PC1 and PC2). Differentiation status was indicated by color. D0: undifferentiated myoblasts, D1-3: 1-3 days after myoblast differentiation began. The percentage of contributing variables for a given PC are shown on the axes. B) Expression levels for ERV-ORFs with ≥ 10 normalized read counts during human primary myoblast differentiation. ERV classes and domain categories are displayed on the left. The regularized logarithm (rlog) transformed read counts for ERV-ORFs are color-coded from blue (low expression) to red (high expression). Differentiation status is presented in the same color as in panel A. C) Human ERV-ORFs overlapping with H3K36me marks. The fractions of samples of histone marks are shown in heat color scales. ERV classes and domain categories on the top are the same as in the panel B. D) Percentages of ERV-ORFs downstream of TSS as obtained from FANTOM datasets in each 1,000 bp bin (all: light blue, transcriptional potential: pink). ERVs without ORFs (non-ERV-ORFs) are shown for comparison (grey). The x-axis represents the distance between ERV-ORFs and the closest TSS. An asterisk (*) indicates statistically significant differences when comparing numbers of the observed ERV-ORF to those expected using fractions of the non-ERV-ORF for each bin (p < 0.001, chi-squared test, FDR corrected). E) PCA plots for TSSs from CAGE datasets, located within 2,000 bp upstream of ERV-ORFs. Colors represent different tissues/cell lines. COBLa_rind: COBL-a (a cell line established from human umbilical cord blood) infected by rinderpest. H9EB/ES, H9 embryoid bodies/embryonic stem cells; MSC, mesenchymal stem cells; RPE, retinal pigment epithelium. F) Percentage of the Repbase annotations exhibiting ERV-ORFs. The different ERV sets for all ERV-ORFs in this study (gray bar), detected from the CHESS datasets (pink circle), histone datasets (yellow circle), Myoblast RNA-Seq datasets (blue circle) are indicated. ERV classes on the left are the same in the panel B. G) Overlaps of human ERV-ORFs with transcription potential detected by different functional datasets. Fractions of ERV-ORFs located near TSSs (< 5kbp) in each functional datasets are presented under the name of functional data.

**Figure 4.**
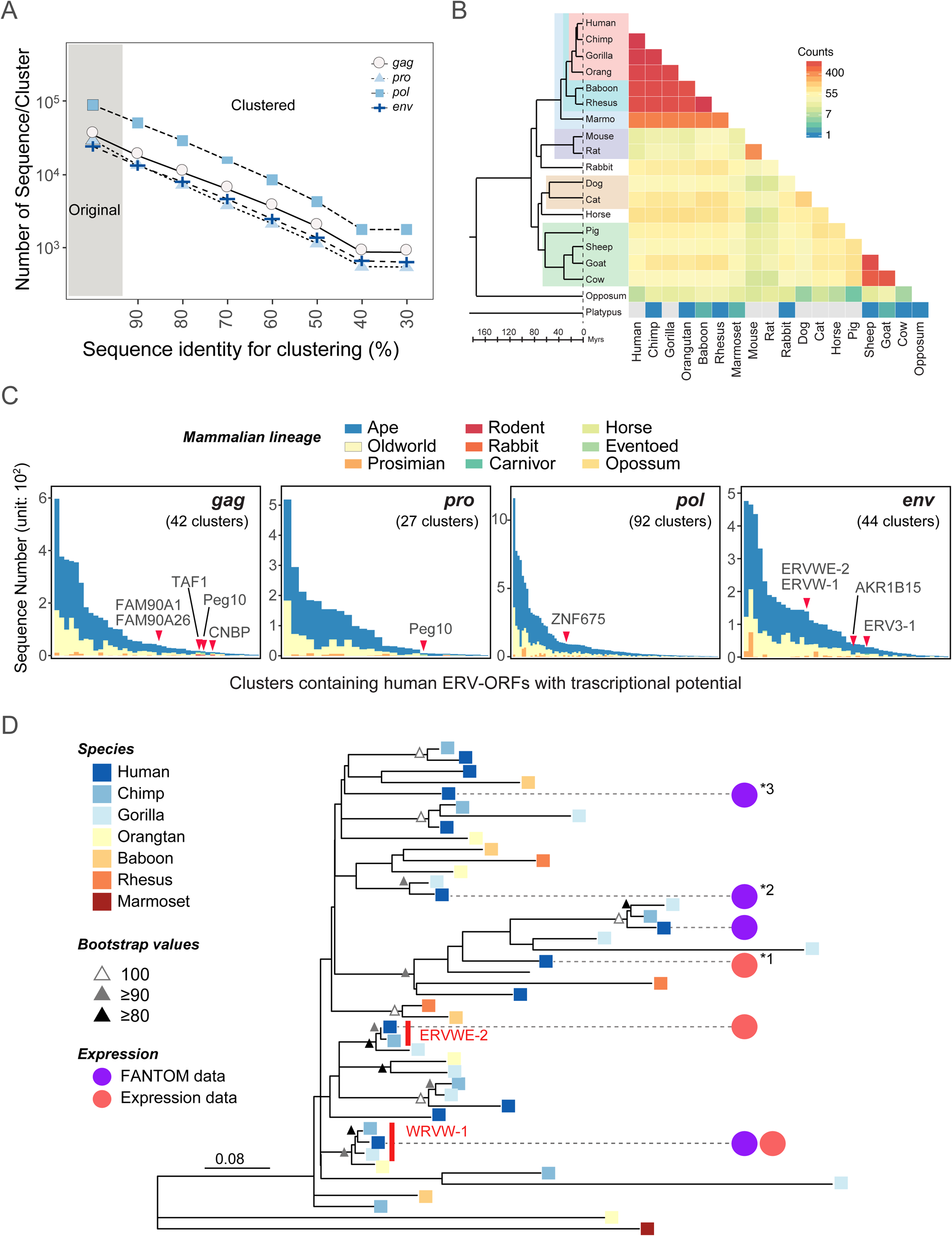
Summary of ERV sequence clustering. A) Number of ERV-ORF clusters associated with each ERV-ORF domain. The x-axis represents the sequence identity established via clustering. The y-axis represents the number of clusters and also shows the original number of sequences before clustering occurred (gray shaded area) for comparison. A logarithmic scale was used on the y-axis. B) Pairwise comparison of shared sequences clustered at ≥ 60% identity level in 19 mammalian species. The color bar represents the number of shared sequences. Gray indicates that no shared sequences were identified. C) The number of ERV-ORF sequences in each cluster containing at least one human ERV-ORF identified in the CHESS database. The x-axis represents clusters with ≥ 60% identity; however, the cluster name is not shown due to limited space. Individual bar colors indicate which ERV-ORFs in each cluster are derived from which species shown in Figure 1A [*e.g*. Apes (blue) contain ERV-ORFs from human, chimpanzee, gorilla, and/or orangutan]. Clusters containing ERV-derived genes are indicated by red triangles and the specific gene name. D) Phylogenetic tree of an ERV-ORF cluster containing six human ERV-ORFs with transcriptional potential. The color of each node represents different ERV-ORF species. Red and pink circles indicate human ERV-ORFs detected in the expression data (CHESS data or myoblast RNA-Seq data) and FANTOM datasets, respectively. The *ERVW-1* and *ERVWE-2* clades are indicated by red font.

One of the features associated with protein-coding genes is the chromatin mark of trimethylated histone H3 at lysine 36 (H3K36me3). We, therefore, applied this feature to identify ERV-ORFs with transcriptional potential as proteins. We obtained the histone mark data of H3K36me3 (9 and 6 cell/tissue types in humans and mice, respectively) from ChIP-Atlas (Oki et al. 2018), and identified the ERV-ORFs overlapping with this mark in a minimum of four samples from each cell/tissue type. From these results we identified 51 and 250 ERV-ORFs overlapping with the mark, in humans and mice, respectively. Further, a higher proportion of samples containing ERV-ORFs with H3K36me3 in each cell/tissue type showed cell type specific patterning (Figure 3B and Figure S6). We also identified several ERV-ORFs marked by multiple cell types; some of which were the fourth exon of CNBP (cellular nucleic acid-binding) genes in humans and mice, yet others were unreported ERV-ORFs. All ERV domains were determined contain the H3K36me3 mark in humans and mice. However, a portion of the domain expression exhibited tissue specificity; for example, in humans, the *pro* domain was found exclusively in liver cells.

To further investigate whether ERV-ORFs are expressed as mRNAs, we examined the presence of transcription start sites (TSS) of ERV-ORFs using the cap analysis of gene expression (CAGE) data in the FANTOM5 database (Lizio et al. 2015). CAGE data are derived from 1,816 human samples from various tissues, primary cells, and cell lines. Using the data, we assessed the relationship between ERV-ORFs located within ten known ERV-derived single-exon genes and the closest TSSs in humans and mice; these ERV-ORFs were located approximately 7,000 bp from the TSS (Table S1). Four ERV-ORFs were located within 1,000 bp of the TSS; however, the rest, specifically those within the *pol* and *env* domains, were found 1,821 bp to 7,086 bp downstream of the TSS (Table S1). Within mice, the ERV-ORFs demonstrated a similar pattern as that observed in humans (Table S1). This suggests that the structure of ERV affects the location at which it integrates into the host genome, and vice versa. We applied this analysis to all our ERV data sets, and further identified ERVs with or without ORFs that were located downstream of the TSS. We calculated the number of ERV-ORFs and non-ERV-ORFs found within each bin of 1,000 bp from the TSS, and found that significantly fewer ERV-ORFs were located within 20,000 bp of TSS compared to non-ERV-ORFs in humans and mice (Figures 3C and S6). By contrast, a significantly larger percentage of ERV-ORFs with transcriptional potential, which were detected by comparing RNA-Seq and histone mark data, were located downstream of TSS compared to Non-ERV-ORFs, especially within 1,000bp in humans [false discovery rate (FDR) corrected p-value < 0.05]. When comparing numbers of the two types of ERV-ORFs within each bin of 2000 bp, we found a significantly larger percentage of ERV-ORFs with transcriptional potential within 4000 bp from TSS compared to non-ERV-ORFs in humans and mice (Figures 3D and S6, FDR corrected p-value < 0.01).

We thus examined which cell types contain active TSSs located upstream of ERV-ORFs in human expression profiles within CAGE data in the FANTOM5 database. We performed PCA analyses using profiles from 38 different cell lines or tissues types, all of which had a minimum of ten available samples for analysis, and found that these active TSSs located near ERV-ORFs were segregated into clearly defined groups for different cell types. Principally, in the expression profile of ERVs, embryonic stem (ES) cells, embryoid body (EB) cells, and induced pluripotent stem (iPS) cells were separated into three groups (Figure 3D). Distinct groups were also identified in the profiles of ERV-ORF in ovary, blood, and brain samples. Specifically, within the ERV profile, osteoblasts established a very clear distinct group.

The Repbase annotations for ERV-ORFs with transcriptional potential were identified as different from those of all other ERV-ORFs in the human genome (Figure 3F). Moreover, the top ten annotations of enriched ERV-ORF were derived from ERV1 and ERVK in humans; while ERV-ORFs with transcriptional potential in three datasets were over-represented in HERVK3, HERVS71, HERV3, HERVFH21 and HERVH48. However, HERV-H was significantly under-expressed in the ERV-ORFs detected in all datasets compared to other ERV-ORFs. Similarly, differences were observed in the Repbase annotations for the ERV-ORFs located within CHESS transcripts and other datasets. For example, high levels of HERV-E and HERVS71 were observed in ERV-ORFs from the CHESS database yet were not observed in ERV-ORFs identified with other datasets. Conversely, HERVK9 and HERVH48 were found to be highly expressed in H3K36me3 histone dataset compared to those detected in CHESS datasets. The histone datasets contain a larger variety of samples, including iPS, ES, and cancer cell lines, while CHESS datasets are obtained exclusively from human tissues. Hence, the observed differences in the annotations of ERV-ORFs may result from differences in sample types used in each project. Indeed, the ERV-ORFs detected via three different functional datasets were quite different (Figure 3G). Although we identified 408 ERV-ORFs with transcriptional potential, only 15 were identified in both the CHESS and histone datasets. Moreover, since myoblast cells were not included in two of the datasets, only 11 ERV-ORFs were found to be overlapped with those in CHESS and histone datasets. These observations confirmed that ERV-ORFs were tissue-specific, and approximately 3.2% (408/12,879) and 2.3% (752/32,062) of ERV-ORFs may be transcribed as proteins in humans and mice, respectively.

### Genome-wide comparison of ERV-ORFs

To understand long-range relationships between mammalian ERVs, we conducted a genome-wide cluster analysis on ERV-ORFs using CD-HIT (Li and Godzik 2006). Since the current annotations by Repbase are based on nucleotide identity, analysis of the amino acid sequences enabled us to better understand the relationship between ERV sequences. Cross-species ERV clustering also provided important predictions for the number of sequence types (different amino acid sequences) in the 19 examined mammalian species, as well as deeper evolutionally relationships. The CD-HIT program clusters sequences by finding the longest reference sequence in each cluster group, and performs subsequent calculations to identify all sequences that contain the reference sequence (Li and Godzik 2006). This clustering step was repeated in a round-robin fashion for each ERV domain category. After clustering ERV-ORF sequences, we obtained 123,035 to 4,092 clusters at the levels of ≥90% to ≥30% sequence identity, respectively (Figure 4A). Furthermore, among ERV genes, the number of clusters identified for *gag* genes (Figure 4A) were more variable, while that for *pro* genes was less variable. We assigned cluster IDs for each cluster by corresponding the number of sequences in the cluster with the length of the reference sequences (Figures S13 and S7).

Cluster analysis identified numerous clusters shared between multiple species, which had not been detected by clustering of nucleotide ERV sequences. Many clusters were shown to be lineage-specific; yet at ≥60% identity, many clusters were found to be shared between primates, rodents, or bovids, whereas less to no, shared clustering occurred in opossum and platypuses (Figure 4B). Using clustering patterns and human RNA-Seq data, we determined the degree of shared ERV-ORFs with transcriptional potential by examining whether the ERV-ORFs were derived from human, primate, or multi-species clusters. Many of the human ERV-ORFs identified in transcripts via RNA-Seq that were detected in transcripts in the CHESS database, and in myoblast differentiation transcriptome data, were also found in clusters of primates (apes and old-world monkeys) at the level of 80%. Furthermore, more than half of the clusters contain only small numbers (< 10) of primate ERVs, indicating they are unique sequences (Figure S8). These human ERV-ORFs with transcriptional potential were also in primate clusters even at the level of ≥60% identity (Figure 4C). Overall, the clustering of *gag* and *env* domains were shared between similar species; many of which were shared among closely related species such as primates, while other *gag* clusters were shared among distantly related species. Additionally, the *pro* domain cluster, containing transcripts found in the CHESS database, was relatively small, and appeared to be more specific to apes compared to other species gene categories. The clusters containing human ERV-ORFs that overlapped with RNA-Seq transcripts with known ERV gene annotations, are highlighted in Figure 4C. Many of these genes, save for syncytin-1, were found in clusters comprised of a smaller number of sequences (≤10). Further, of the total 12,879 human ERV-ORFs examined in this study, only 223 sequences were human-specific (un-clustered unique sequences or human-specific clusters). Moreover, from the ERV-ORFs that overlapped with RNA-Seq data transcripts, only two sequences were found to be un-clustered. Mouse ERV-ORFs showed a similar tendency, although the species specificity was stronger than humans. For example, 18,730 sequences out of the total 32,063 ERV-ORFs were mouse-specific, and 349 out of 586 ERV-ORFs detected in the RNA-Seq data were determined to be in mouse-specific clusters. Only 15 ERV-ORFs were unique sequences in the mouse genome.

To understand the relationship between sequence similarity and functionality in each cluster, we analyzed one cluster containing six human ERV-ORFs with transcriptional potential. We extracted 44 unique mammalian ERV-ORF sequences with the lowest detectable similarity (≥ 30%), that were ≥250 aa in length from the cluster (Figure 4C, *env* panel), and generated a maximum likelihood phylogeny by using the amino acid sequences using RAxML (Stamatakis 2006) (Figure 4D). Our results determined that this is a primate-specific cluster and contains *ERVW-1*, which encodes syncytin-1. Syncytin-1 has been found only in apes, and thus *ERVW-1* formed a small clade with apes, not with monkeys (Figure 4D). Additionally, the branch length for apes within the syncytin-1 clade was shorter compared to other clades, which may indicate evolutional conservation of sequences. We also found another small clades containing short branch lengths among apes. The human ERV-ORF in this clade was identified in the RNA-Seq myoblast differentiation (Figure 4D). The ERV-ORF was overlapped with *ERVWE-2* (endogenous retrovirus group W member). Note that the other ERV-ORF overlapping with CHESS transcripts did not contain clades at this specific cut-off (Figure 4D); however, it did have a similar ERV-ORF in chimpanzees when the length cut-off was relaxed (the length of chimpanzee ERV-ORF was 229 aa), and exhibited a short branch length. In addition, we examined the relationship between the human ERV-ORFs in the phylogeny and TSSs using FANTOM5 datasets, and found that four human ERV-ORFs were located 2,000 bp downstream of the TSSs (Figure 4D, purple circle). Although other human ERV-ORFs in the tree were not found in our functional datasets, two sequences that were detected exclusively with the TSS data were determined to be perfectly aligned with unannotated ESTs (Figure 4D). The transcript overlapping with the former ERV-ORF was found in brain, skeletal muscle, and cancer cells (Yi et al. 2004; Schmitt et al. 2013), with the transcript overlapping with the latter found in placental tissues (Dias Neto et al. 2000).

Many human ERV-ORFs identified via RNA-Seq analysis were found to be shared among primates, while some ERV-ORFs were found in multiple shared clusters. Indeed, of the clusters at the level of 60% identity, we identified 60 clusters that were shared among ≥8 mammalian species (Figure 5). The largest shared ERV-ORF domain category was for the *pol* gene, which had 27 clusters shared between species. The remaining ERV gene domains contained a similar number of cross-species clusters (12, 11, and 10 for *gag, pro*, and *env*, respectively). Although nine clusters contained known annotated genes or reported transcripts (Figure 5 and Table S1), many of the ERV-ORFs identified in the clusters were previously unreported; seven clusters contain human ERV-ORFs with transcriptional potential but not known ERV-derived genes. No clusters containing only mouse ERV-ORF with transcriptional potential was found. Interestingly, 15 of these multi-species clusters did not contain human ERV-ORFs.

**Figure 5.**
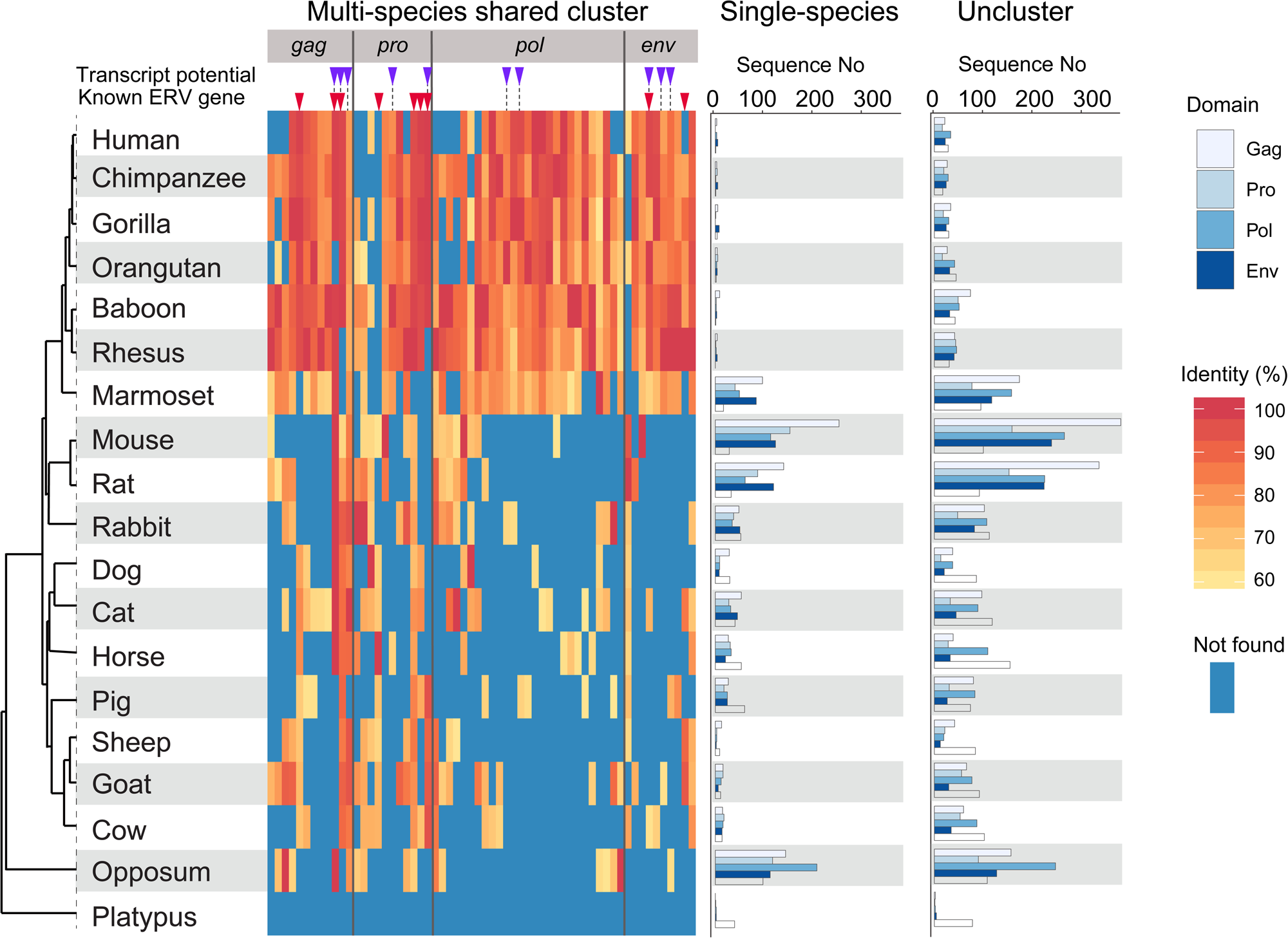
ERV-ORF clusters shared among at least eight species. Clusters of ERV-ORF sequences shared among mammals at ≥ 60% identity levels are shown. Domain names are shown on the top. Clusters containing ERV-ORFs with transcriptional potential (purple triangle) and known genes (red triangle) are highlighted. The red-scale color bar represents the percent identities for sequences against a reference sequence within the cluster. The highest identity in each species is shown. Blue represents the absence of a specific sequence in the given cluster. The clusters containing human ERV-ORFs with transcriptional potential (purple triangle) and known mammalian ERV genes (red triangle) were indicated on the top of heatmap. The amount of sequences forming single-species clusters (middle) and those that failed to form any cluster (right) are also shown for each species.

It is also worthy to note that nine of these clusters remained following the extraction of ≥8 shared species clusters at the level of ≥80% identity, and 21 clusters remained when extracting clusters shared among ≥10 mammalian species (Figures S9A and S9B). Further, there were fewer multi-species ERV-ORF clusters identified in rodents and opossum. Instead, these species exhibited larger numbers of species-specific clusters and unique ERV sequences (Figure 5, right panels). The species-specific clusters in these species were abundant in *pol* genes, while there were no multi-species shared clusters observed in analysis of platypuses.

## Discussions

We comprehensively identified ERV-ORFs in 19 mammalian species, with differences in their domain fractions noted between species (Figures 1A and 1B). This is the first report, to our knowledge, that presents data on the genome-wide prediction of ERV-ORFs at the domain level and their expressions in a wide range of mammalian species. In general, ERV-ORFs are not considered as protein-coding genes. However, our results clearly indicate that approximately 2-3% of ERV-ORFs may be transcriptionally active in humans and mice, respectively. Although the percentage of ERV-ORFs that function as potential candidates for functional genes was determined to be small, there may be unidentified protein-coding genes in these ERV-ORFs.

Alternatively, the observed ERV-ORFs may have been present by chance after recently becoming inserted into the host genome; as was observed with an endogenous bornavirus-like nucleoprotein element (EBLNs) expressed in simians (Kobayashi et al. 2011). The observed positive correlation between the fraction of ERV-ORFs and the total ERV length may reflect this. To exclude ERV-ORFs that were simply present by chance without a known function, we compared the divergence distribution of our observed ERV-ORFs to those presented by Repbase consensus sequences using human ERV-ORFs, which are considered to exhibit high transcription potential. Note that ERV-ORFs identified by functional data included those located ≤ 2,000 bp downstream of the TSSs. When comparing divergence patterns, we observed an interesting change in the frequency of ERV-ORFs with transcriptional potential. At a divergence of approximately 4-7%, the frequency of ERV-ORFs with transcriptional potential in humans was determined to be lower than the total number of ERV-ORFs (Figures S10). This confirmed that ERV-ORFs with relatively small divergence (approximately ≤ 7%), were likely recently inserted into the genomes, and may include ERV-ORFs that remain by chance alone.

We also noted that entire ERV-ORFs were significantly depleted near the TSS in humans and mice. This may support the low transcriptional potential of ERV-ORFs as a whole. If specific epigenetic changes occur in close proximity to ERV-ORFs with low transcriptional potential, they subsequently begin to be expressed, which may lead to development of diseases as described in previous studies (Perron et al. 2012; Kassiotis 2014). This may provide an explanation for why ERV-ORFs become depleted close to TSS in transcriptionally active regions, to avoid potential detrimental secondary effects to their expression. Hence, these preserved ERV-ORFs in transcriptionally inactive regions may function as the raw materials required for constructing new exons for multi-exon genes composed of non-ERV-derived exons during mammalian evolution. Indeed, we identified ERV-ORFs that were exonized in known Ensembl multi-exon genes. Further, the numbers of ERV-containing genes varied among mammalian species (Figure S5); horse, sheep, and opossum contained a larger number of these genes compared to other mammalian species. It is unclear whether this observation reflected the difference in ERV-ORF co-opted rates or simply in gene annotation quality. Nevertheless, differences, although small, were observed in the number of ERV-ORFs identified in well-annotated human and mouse genomes (Figure S5). We also found that certain ERV-ORFs partially overlapped with known gene exons (e.g. *AKR1B15, UBXN8*, and *PPHLN1* in humans) containing splice variants. This suggests that, as reported in a previous study (Bae et al. 2013), ERV-ORFs may be transcribed as alternative transcript variants. It is interesting to note that known multi-exon genes have been identified as containing ERV-ORFs as predicted by HMM alone, suggesting that these ORFs may be originated from ERVs. Moreover, similar numbers of these genes were identified in all examined mammals, save for horses, sheep, and platypuses. Considering that orthologs of these genes were found in different species (e.g. *CNBP, GIN1, SUGP2*, in chickens and *CTSE* in humans and mice), these ERV-ORFs may have been co-opted before the evolutionary divergence of birds and mammals.

The ERV-ORF expression results may also suggest that the promoter regions of ERVs were co-opted rather than the ORFs. Previous studies have reported that promoter regions, most notably for ERV LTRs, have been co-opted as tissue-specific, or alternative promoters in mammalian cells (well summarized in Thompson et al. 2016). In this analysis, we were unable to exclude this as a possibility. The LTR promoter usage in a given cell may also be inferred from RNA-Seq data. When analyzing promoter usage, the expression levels were often accumulated and applied to each ERV group, such as HERV-K and MERV-L. Thus, if only a certain ERV-ORF is functional, this accumulated data may cause underestimation of the actual ORF expression levels. In our analysis of human myoblast RNA-Seq data, we presented the expression of each ERV-ORF locus separately, and confirmed that the expression of each ERV-ORF loci was unique during muscle cell differentiation (Figure 3B). We believe this level of detailed expression analysis will assist in revealing the function of ERV-ORFs by identifying individual ERV-ORF loci.

Combinatorial analysis of functional datasets and cluster analysis for ERV-ORFs in humans and mice revealed that many expressed ERV-ORFs were tissue- and lineage-specific. This is consistent with previous studies showing that specific ERV genes, such as *ERVW-1,* are expressed in limited tissues and are shared exclusively among apes. In addition, we demonstrated that fractions of each viral-like protein domain in ERV-ORFs varied among mammalian lineages. Further, ERV-ORF fractions containing transcriptional potential were not significantly different from those of entire ERV-ORFs in humans and mice. Moreover, we observed that the fractions of viral-like protein domains among mammalian linages were quite different. This implies that the fractions of retroviral-like domains within ERV-derived genes also differ among mammalian lineages, and are differentially transcribed in different tissues, which may affect lineage-specific characteristics at varying levels.

Together, our results suggest that mammalian ERV-ORFs, many of which were not previously described, may be co-opted in a lineage-specific manner, and are likely to have contributed to mammalian evolution and diversification. However, functional data is currently very limited. Thus, further ERV-ORF studies employing the functional data from multiple species may provide a deeper understanding of mammalian evolution.

## Materials and methods

### Extraction of ERV-ORFs

All sequences and annotations for potential protein-coding ERV-ORFs from 19 mammalian genomes were used in this study were obtained from the gEVE database (Nakagawa and Takahashi 2016; http://geve.med.u-tokai.ac.jp). Species names and corresponding genome versions used in this study are summarized in Table S2. We classified ERV-ORF sequences by domains predicted by HMMER version 3.1b1 (hmmer.org) with HMM profiles and/or RetroTector (Sperber et al. 2007) as provided in the gEVE database. We further separated ERV *pol* and LINE-ORF2 domains as previously described (Nakagawa and Takahashi 2016). We first calculated the fraction of ERV-ORF domains within a species, and obtained the average for each domain across all 19 mammals. We then further divided the original domain fractions for each species by the mammalian average and obtained fold changes for all domains.

### Divergence of ERV-ORFs

We used RepeatMasker (Smith 2013) to calculate CpG adjusted divergence information [substitutions levels calculated by Kimura-two parameter (Kimura 1980) for ERV-ORFs]. The R package version 3.4.4 (R Core Team 2017) was used to calculate the means for each specie’s genome. We determined the modes of divergence using the R package LaplacesDemon version 16.1.1 (Statisticat 2018).

### RNA-Seq analysis

RNA-Seq data for human and mouse myoblast differentiation were downloaded from the NCBI SRA; https://www.ncbi.nlm.nih.gov/sra). For humans, we used 12 RNA-Seq datasets from skeletal muscle cells under the SRA study number SRP033135 (Trapnell et al. 2014). We obtained 14 RNA-Seq datasets from C2C12 cell differentiation under the SRA number SRP036149 to analyze mouse myoblast differentiation. The accession number of runs used in this analysis are summarized in Table S3. After trimming the sequences using fastp version 0.12.5 (Chen et al. 2018) with the following set parameters: -q 20 -l 30, reads were mapped onto the human (GRCh38), and mouse (mm10) genomes using HISAT2 version 2.1.0 (Kim et al. 2015) with the default option. StringTie version 1.3.4b (Pertea et al. 2015) and DESeq2 version 1.18.1 (Love et al. 2014) were used to quantify and evaluate EVE expression. In this analysis, we performed mapping and quantification for ERV-ORF transcripts by using in-house ERV gene transfer format (GTF) files, which were available in the gEVE database. After normalizing for size and filtering the data (removing small read counts < 10 aa), we log transformed the count data using the rlog transformation function of the DESeq2, which minimizes the detection of sample differences for transcripts with small counts, and normalizes the counts with respect to the library size. We then performed PCA analysis using the standardized read counts. PCA plots were generated using the built-in R function prcomp. For human RNA-Seq data, expression data was sorted based on the total sum of each ERV-ORF expression profile, and the extracted top 100 ERV-ORF transcripts. The standardized counts were then represented in a heatmap, which was generated using the R package pheatmap version 1.0.10 (Kolde 2015).

### Analysis using CHESS

To investigate the functionality of ERV-ORF, we obtained RNA-Seq datasets from the GTEx project (Carithers et al. 2015) in GFF format from the CHESS v.2.1 database (Pertea et al. 2018). We then identified overlapping ERV-ORFs with exonic regions of transcripts in the GFF file using the intersect function of BEDTools version 2.26.0 (Quinlan et al. 2010).

### Analysis using H3K36me3 histone mark

We obtained histone mark data for H3K36me3 from the Peak Browser in ChIP-atlas (Oki et al. 2018) with threshold for significance set at 50. To extract mark positions with strong evidence, we used only those marks that were detected in a minimum of five samples. Further, for comparison of H3K36me3 marked ERV-ORFs among cell types, we only used the histone marks obtained from more than ten samples in each tissue or cell.

### Analysis using FANTOM data

To predict which ERV-ORFs become transcribed, we analyzed CAGE data from FANTOM5 (Lizio et al. 2015). Human and mouse CAGE data (bed files, read count data, and sample information table) were retrieved from the FANTOM5 data repository. We converted ERV-ORF coordinates into hg19 assembly using the UCSC LiftOver tool (Hinrichs et al. 2006), and identified the ERV-ORF and non-ERV-ORF located downstream of TSSs using the closest function in BEDTools version 2.26.0. To determine which of the differences in number of ERV-ORF and non-ERV-ORFs near TSS were significant, Fisher’s exact test was performed using built-in R function. The expected number of ERV-ORF located downstream of TSSs with a bin size of 1000 bp and 2000 bp was estimated using the fraction of non-ERV-ORF with the same bin size. To control for the FDR, we performed multiple-test correction with FDR (Benjamini and Hochberg 1995) using the p.adjust function in the R package. PCA analysis for TSS data was performed using the same procedure as described for the RNA-Seq data.

### Cluster analysis

We obtained ERV-ORF alignments, and clustered each domain with the CD-HIT program (Li and Godzik, 2006) for 40-90% shared sequence identity, and we used the PSI-CD-HIT program (Altschul et al. 1997) to detect ≤ 30% level identity. We then extracted the sequence that contained the highest level of homology with the reference sequence for each species within each cluster and generated matrices for heatmaps. Manipulation of this data was accomplished using in-house Perl scripts. Heatmaps was visualized using the geom_tile function of the R package ggplot2 (Wickham 2016).

### Phylogenetic analysis

We first identified clusters that contained six human ERV-ORFs including *ERVW-1* in the *env* domain with ≥ 30% identity homology, and extracted amino acid sequences that were ≥ 250 aa in length from that cluster. After multiple alignments of the sequences with MAFFT version 7.394 using the auto option (Katoh and Standley 2013), we generated maximum likelihood trees using RAxML version 8.2.11 with the following options: -f a -m PROTGAMMAJTTF -#500; 50. The phylogenetic tree was visualized using the R package ggtree version 1.10.5 (Yu et al. 2017).

### Data manipulations and visualizations

For all analyses employed in this study, we used in-house programs developed with Perl, Python, AWK, and Shell script as well as R package dplyr (Wickham et al. 2018) and rehape (Wickmham 2007) for processing and manipulation of the data. For visualization, we used ggplot2 version 2.2.1 with RColorBrewer (Wickham 2016) in the R package.

## List of abbreviations

ERV: endogenous retroviruses
ERV-ORF: endogenous retrovirus open reading frame
TSS: transcription start site
TE: transposable element
HMM: hidden Markov model
CAGE: cap analysis of gene expression
PCA: Principal component analysis
GTF: gene transfer format
SRA: sequence read archive
GTEx: genotype-tissue expression
CHESS: comprehensive human expressed sequence
ES: embryonic stem
EB: embryoid body
iPS: induced pluripotent stem

## Acknowledgments

This work was supported by JSPS KAKENHI Grants-in-Aid for Challenging Exploratory Research (17K19359 to MTU and SM), Young Scientists (16K21386 to SN), Scientific Research on Innovative Areas (16H06429, 16K21723, 17H05823, 19H04843 to SN) and by MEXT-Supported program for the Strategic Research Foundation at Private Universities (S1411010 to MTU and SN). Computations were performed in part on the NIG supercomputer at ROIS National Institute of Genetics and SHIROKANE at Human Genome Center (the Univ. of Tokyo).

## References

Altschul SF, Madden TL, Schaff er AA, Zhang J, Zhang Z, Miller W, Lipman DJ. 1997. Gapped BLAST and PSI-BLAST: a new generation of protein database search programs. Nucleic Acids Res. 25:3389–3402.

Ashley J, Cordy B, Lucia D, Fradkin LG, Budnik V, Thomson T. 2018. Retrovirus-like Gag Protein Arc1 Binds RNA and Traffics across Synaptic Boutons. Cell 172:262–274.

Bae MI, Kim YJ, Lee JR, Jung YD, Kim HS. 2013. A new exon derived from a mammalian apparent LTR retrotransposon of the SUPT16H gene. Int J Genomics 2013:387594.

Bannert N, Kurth R. 2006. The evolutionary dynamics of human endogenous retroviral families. Annu Rev Genomics Hum Genet. 7:149–173.

Bao W, Kojima KK, Kohany O. 2015. Repbase Update, a database of repetitive elements in eukaryotic genomes. Mob DNA 6:11.

Bénit L, Calteau A, Heidmann T. 2003. Characterization of the low-copy HERV-Fc family: evidence for recent integrations in primates of elements with coding envelope genes. Virology 312:159–168.

Benjamini, Y. Hochberg Y. 1995. Controlling the False Discovery Rate: A Practical and Powerful Approach to Multiple Testing. J R Stat Soc Ser A Stat Soc. 57: 289–300.

Bolze PA, Mommert M, Mallet F. 2017. Contribution of syncytins and other endogenous retroviral envelopes to human placenta pathologies. Prog Mol Biol Transl Sci. 145: 111–162.

Carithers LJ, Ardlie K, Barcus M, Branton PA, Britton A, Buia SA, Compton CC, DeLuca DS, Peter-Demchok J, Gelfand ET, et al. 2015. A Novel Approach to High-Quality Postmortem Tissue Procurement: The GTEx Project. Biopreserv Biobank 13:311–319.

Chen S, Zhou Y, Chen Y, Gu J. 2018. fastp: an ultra-fast all-in-one FASTQ preprocessor. Bioinformatics 34: i884–i890.

Chuong EB, Elde NC, Feschotte C. 2016. Regulatory evolution of innate immunity through co-option of endogenous retroviruses. Science 351:1083–1087.

de Koning AP, Gu W, Castoe TA, Batzer MA, Pollock DD. 2011. Repetitive elements may comprise over two-thirds of the human genome. PLoS Genet. 7:e1002384.

de Parseval N, Lazar V, Casella JF, Benit L, Heidmann T. 2003. Survey of human genes of retroviral origin: Identification and transcriptome of the genes with coding capacity for complete envelope proteins. J Virol. 77:10414–10422.

Dias Neto E, Correa RG, Verjovski-Almeida S, Briones MR, Nagai MA, da Silva W Jr, Zago MA, Bordin S, Costa FF, Goldman GH, et al. 2000. Shotgun sequencing of the human transcriptome with ORF expressed sequence tags. Proc Natl Acad Sci U S A. 97:3491–3496.

Diwash J, Feschotte C, Betran E. 2017. Transposable Element Domestication As an Adaptation to Evolutionary Conflicts. Trends Genet. 33: 817–831.

Dupressoir A, Lavialle C, Heidmann T. 2012. From ancestral infectious retroviruses to bona fide cellular genes: role of the captured syncytins in placentation. Placenta 33, 663–667.

Grow EJ, Flynn RA, Chavez SL, Bayless NL, Wossidlo M, Wesche DJ, Martin L, Ware CB, Blish CA, Chang HY, et al. 2015. Intrinsic retroviral reactivation in human preimplantation embryos and pluripotent cells. Nature 522:221–225.

Hinrichs, AS, Karolchik D, Baertsch R, Barber GP, Bejerano G, Clawson H, Diekhans M, Furey TS, Harte RA, Hsu F, et al. 2006 The ucsc genome browser database: update 2006. Nucleic Acids Res. 34(Suppl 1):D590–D598.

International Human Genome Sequencing Consortium. 2001. Initial sequencing and analysis of the human genome. Nature 409:860–921.

Ito J. Sugimoto R. Nakaoka H, Yamada S, Kimura T, Hayano T, Inoue. 2017. Systematic identification and characterization of regulatory elements derived from human endogenous retroviruses. PLoS Genet. 13: e1006883.

Kassiotis G. 2014. Endogenous retroviruses and the development of cancer. J Immunol. 192:1343–1349.

Katoh K, Standley DM. 2013. MAFFT multiple sequence alignment software version 7: improvements in performance and usability. Mol Biol Evol. 30: 772–780.

Kim D, Langmead B, Salzberg SL. 2015. HISAT: a fast spliced aligner with low memory requirements. Nat Methods 12:357–360.

Kimura M. 1980. A simple method for estimating evolutionary rates of base substitutions through comparative studies of nucleotide sequences. J Mol Evol. 16:111–120.

Kobayashi Y, Horie M, Tomonaga K, Suzuki Y. 2011. No evidence for natural selection on endogenous borna-like nucleoprotein elements after the divergence of Old World and New World monkeys. PLoS One 6:e24403.

Kolde R. 2015. pheatmap: Pretty Heatmaps. R package version 1.0.8. https://CRAN.R-project.org/package=pheatmap

Lavialle C, Cornelis G, Dupressoir A, Esnault C, Heidmann O, Vernochet C, Heidmann T. 2013. Paleovirology of ‘syncytins’, retroviral env genes exapted for a role in placentation. Philos Trans R Soc Lond B. 368:20120507.

Li W, Godzik A. 2006. Cd-hit: a fast program for clustering and comparing large sets of protein or nucleotide sequences. Bioinformatics 22:1658–1659.

Lin L, Xu B, Rote NS. 1999. Expression of endogenous retrovirus ERV-3 induces differentiation in BeWo, a choriocarcinoma model of human placental trophoblast. Placenta 20:109–118.

Lizio M, Harshbarger J, Shimoji H, Severin J, Kasukawa T, Sahin S, Abugessaisa I, Fukuda S, Hori F, Ishikawa-Kato S, et al. 2015. Gateways to the FANTOM5 promoter level mammalian expression atlas. Genome Biol. 16:22.

Love MI, Huber W, Anders S. 2014. Moderated estimation of fold change and dispersion for RNA-seq data with DESeq2. Genome Biol. 15:1–21.

Matsui T, Miyamoto K, Kubo A, Kawasaki H, Ebihara T, Hata K, Tanahashi S, Ichinose S, Imoto I, Inazawa J, et al. 2011. SASPase regulates stratum corneum hydration through profilaggrin-to-filaggrin processing. EMBO Mol Med. 3:320–333.

McVean G. 2010. What drives recombination hotspots to repeat DNA in humans? Phillos Trans R Soc B Biol Sci. 365, 1213–1218.

Mi S, Lee X, Li X, Veldman GM, Finnerty H, Racie L, LaVallie E, Tang XY, Edouard P, Howes S, et al. 2000. Syncytin is a captive retroviral envelope protein involved in human placental morphogenesis, Nature 403: 785–789.

Mouse Genome Sequencing Consortium. 2002. Initial sequencing and comparative analysis of the mouse genome. Nature 420:520–562.

Nakagawa S, Takahashi MU. 2016. gEVE: a genome-based endogenous viral element database provides comprehensive viral protein-coding sequences in mammalian genomes. Database 2016:baw087.

Nakaya Y, Koshi K, Nakagawa S, Hashizume K, Miyazawa T. 2013. Fematrin-1 is involved in fetomaternal cell-to-cell fusion in Bovinae placenta and has contributed to diversity of ruminant placentation. J Virol. 87:10563–10572.

Neuwirth E. 2014. RColorBrewer: ColorBrewer Palettes. R package version 1.1-2. https://CRAN.R-project.org/package=RColorBrewer

Nishihara H, Kobayashi N, Kimura-Yoshida C, Yan K, Bormuth O, Ding Q, Nakanishi A, Sasaki T, Hirakawa M, Sumiyama K, et al. 2016. Coordinately co-opted multiple transposable elements constitute an enhancer for wnt5a expression in the mammalian secondary palate. PLoS Genet. 12:e1006380.

Oki S, Ohta T, Shioi G, Hatanaka H, Ogasawara O, Okuda Y, Kawaji H, Nakaki R, Sese J, Meno C. 2018. ChIP-Atlas: a data-mining suite powered by full integration of public ChIP-seq data. EMBO Rep. 19: e46255.

Ono R, Nakamura K, Inoue K, Naruse M, Usami T, Wakisaka-Saito N, Hino T, Suzuki-Migishima R, Ogonuki N, Miki H, et al. 2006. Deletion of Peg10, an imprinted gene acquired from a retrotransposon, causes early embryonic lethality, Nat Genet. 38: 101–106.

Paces J, Pavlícek A., Paces V. 2002. HERVd: database of human endogenous retroviruses. Nucleic Acids Res. 30:205–206.

Pastuzyn ED, Day CE, Kearns RB, Kyrke-Smith M, Taibi AV, McCormick J, Yoder N, Belnap DM, Erlendsson S, Morado DR, et al. 2018. The Neuronal Gene Arc Encodes a Repurposed Retrotransposon Gag Protein that Mediates Intercellular RNA Transfer. Cell 172:275–288.e18.

Perron H, Germi R, Bernard C, Garcia-Montojo M, Deluen C, Farinelli L, Faucard R, Veas F, Stefas I, Fabriek BO, et al. 2012. Human endogenous retrovirus type W envelope expression in blood and brain cells provides new insights into multiple sclerosis disease. Mult Scler. 18:1721–1736.

Pertea M, Pertea GM, Antonescu CM, Chang TC, Mendell JT, Salzberg SL. 2015. StringTie enables improved reconstruction of a transcriptome from RNA-seq reads. Nat Biotechnol. 33:290–295.

Pertea M, Shumate A, Pertea G, Varabyou A, Breitwieser FP, Chang Y, Madugundu AK, Pandey A, Salzberg SL. 2018. CHESS: a new human gene catalog curated from thousands of large-scale RNA sequencing experiments reveals extensive transcriptional noise. Genome Biol. 19:208.

Quinlan AR, Hall IM. 2010. BEDTools: a flexible suite of utilities for comparing genomic features. Bioinformatics 26:841–842.

R Core Team. 2017. R: A language and environment for statistical computing. R Foundation for Statistical Computing, Vienna, Austria. https://www.R-project.org/

Schmitt K, Richter C, Backes C, Meese E, Ruprecht K, Mayer J. 2013. Comprehensive analysis of human endogenous retrovirus group HERV-W locus transcription in multiple sclerosis brain lesions by high-throughput amplicon sequencing. J Virol. 87:13837–52.

Sela N, Mersch B, Gal-Mark N, Lev-Maor G, Hotz-Wagenblatt A, Ast G. 2007. Comparative analysis of transposed element insertion within human and mouse genomes reveals Alu’s unique role in shaping the human transcriptome. Genome Biol. 8:R127.

Smit AFA. 1999. Interspersed repeats and other mementos of transposable elements in mammalian genomes. Curr Opin Genet Develop. 9: 657–663.

Smit AFA, Hubley R, Green P. 2013. RepeatMasker Open-4.0., 2013-2015. http://www.repeatmasker.org.

Sperber GO, Airola T, Jern P, Blomberg J. 2007. Automated recognition of retroviral sequences in genomic data–RetroTector. Nucleic Acids Res. 35:4964–4976.

Stamatakis A. 2006. RAxML-VI-HPC: maximum likelihood-based phylogenetic analyses with thousands of taxa and mixed models. Bioinformatics 22: 2688–2690.

Statisticat LLC. 2018. LaplacesDemon: Complete Environment for Bayesian Inference. Bayesian-Inference.com. R package version 16.1.1. https://web.archive.org/web/20150206004624/ http://www.bayesian-inference.com/software

Subramanian R, Wildschutte J, Russo C, Coffin J. 2011. Identification, characterization, and comparative genomic distribution of the HERV-K (HML-2) group of human endogenous retroviruses. Retrovirology 8:90.

Sugimoto J. Sugimoto M. Bernstein H, Jinno Y, Schust D. 2013. A novel human endogenous retroviral protein inhibits cell-cell fusion. Sci Rep. 3:1462.

Thompson PJ, Macfarlan TS, Lorincz MC. 2016. Long Terminal Repeats: From Parasitic Elements to Building Blocks of the Transcriptional Regulatory Repertoire. Mol Cell 62:766–76.

Thornburg BG, Gotea V, Makalowski W. 2006. Transposable elements as a significant source of transcription regulating signals. Gene 365, 104–110.

Tokuyama M, Kong Y, Song E, Jayewickreme T, Kang I, Iwasaki A. 2018. ERVmap analysis reveals genome-wide transcription of human endogenous retroviruses. Proc Natl Acad Sci USA. 115:12565–12572.

Trapnell C, Cacchiarelli D, Grimsby J, Pokharel P, Li S, Morse M, Lennon NJ, Livak KJ, Mikkelsen TS, Rinn JL. 2014. The dynamics and regulators of cell fate decisions are revealed by pseudotemporal ordering of single cells. Nat Biotechnol. 32:381–386.

Villesen P, Aagaard L, Wiuf C, Pedersen FS. 2004. Identification of endogenous retroviral reading frames in the human genome. Retrovirology 1:32.

Wickham H. 2007. Reshaping Data with the reshape Package. J Stat Softw 21:1–20. http://www.jstatsoft.org/v21/i12/

Wickham H. 2016. ggplot2: Elegant Graphics for Data Analysis. Springer-Verlag New York.

Wickham H, François R, Henry L, Müller K. 2018. dplyr: A Grammar of Data Manipulation. R package version 0.7.5. https://CRAN.R-project.org/package=dplyr

Yi JM, Kim HM, Kim HS. 2004. Expression of the human endogenous retrovirus HERV-W family in various human tissues and cancer cells. J Gen Virol. 5:1203–1210.

Yu G, David Smith, Zhu H, Guan U, Lam TT. 2017. ggtree: an R package for visualization and annotation of phylogenetic trees with their covariates and other associated data. Methods Ecol Evol. 8:28–36.

