## Supplementary table for "Comprehensive genomic analysis reveals dynamic evolution of endogenous retroviruses that code for retroviral-like protein domains"

**Table S1** EVE-ORFs overlapping with Ensembl annotated gene.

**ID**: gEVE database ID, **HMM**: viral motives predicted by HMM, **Repbase**: Repbase annotation of EVE-ORFs predicted by RepeatMasker, **Gene**: gene name and Ensembl transcript or gene IDs. We used transcript ID when available. If the transcript has no gene name, "NA" is shown. ERV-ORFs residing within "known ERV-derived single-exon genes" are indicated by cells colored pink. **Dist_TSS**: the distance to the TSSs is shown only for the ERV-ORFs in the known ERV-derived single-exon genes in humans and mice.

#### Human EVE-ORFs

Only human has a column for CHESS annotations (**CHESS**). In the column, EVE-ORFs overlapping status with CHESS transcript is shown; each EVE-ORF has ≥50% overlap, ≥30%, and ≥10% with CHESS transcript, respectively.

| ID | Domain | HMM | Repbase | Gene | CHESS | Dist_TSS |
|---|---|---|---|---|---|---|
| Hsap38.chr2.69960660.69961718.− | Pro | Pro:AP;AP_DTG_ILG_template\|AP;AP_saspa | . | ASPRV1;ENSP00000315383 | | 0 |
| Hsap38.chr3.129171078.129171320.− | Gag | GAG;GAG_lentiviridae | . | PPHP;ENSP0000422110 | ≥50% | |
| Hsap38.chr4.9173574.9173990.+ | Gag | gag;zf−CCHC_6 | . | FAM90A26;ENSP00000421131 | | |
| Hsap38.chr5.103087901.103088227.− | Pro | INT;GIN1 | . | GIN1;ENSP00000427162 | ≥50% | |
| Hsap38.chr5.88190260.88190814.− | Pol | RNaseH;RNaseH_gammaretroviridae | HERVE−int;LTR/ERV1\|L1MEg;LINE/L1 | TMEM161B;ENSP00000426354 | | |
| Hsap38.chr6.11103697.11105316.− | Env | ENV;ENV_deltaretroviridae\|ENV;ENV_D−type_betaretroviridae\|ENV;ENV_gammaretroviridae\|ENV;ENV_retroviridae | MER50−int;LTR/ERV1 | ERVFRD−1;ENSP00000444461 | | 6408 |
| Hsap38.chr7.134564320.134564778.+ | Env | env;TLV_coat\|ENV;ENV_D−type_betaretroviridae\|ENV;ENV_gammaretroviridae | HERVI−int;LTR/ERV1 | AKR1B15;ENSP00000397009 | ≥50% | 15405 |
| Hsap38.chr7.64991215.64993059.− | Env | ENV;ENV_gammaretroviridae | HERV3−int;LTR/ERV1 | ERV3−1;ENSP00000391594 | | 6410 |
| Hsap38.chr7.92468768.92470387.− | Env | env;TLV_coat\|ENV;ENV_D−type_betaretroviridae\|ENV;ENV_gammaretroviridae | HERV17−int;LTR/ERV1 | ERVW−1;ENSP00000419945 | | 7086 |
| Hsap38.chr7.94663299.94664531.+ | Gag | GAG;GAG_v_clade | (CCT)n;Simple_repeat\|(CAA)n;Simple_repeat | PEG10;ENSP00000480676 | | 0 |
| Hsap38.chr7.94664474.94665679.+ | Pro | AP;AP_v_clade | (ACC)n;Simple_repeat\|(CCG)n;Simple_repeat | PEG10;ENSP00000480676 | | 0 |
| Hsap38.chr8.30731656.30732471.+ | | pol;RVT_1\|RT;RT_17_6\|RT;RT_alpharetroviridae\|RT;RT_betaretroviridae\|RT;RT_deltaretroviridae\|RT;RT_epsilonretroviridae\|RT;RT_gammaretroviridae\|RT;RT_spumaretroviridae | HERVE−int;LTR/ERV1 | UBXN8;ENSP00000477532 | ≥50% | |
| Hsap38.chr8.30732333.30732620.+ | Pol | RT;RT_gammaretroviridae | HERVE−int;LTR/ERV1 | UBXN8;ENSP00000477532 | ≥30% | 4638 |
| Hsap38.chr8.30732669.30733337.+ | Pol | pol;RNase_H\|RNaseH;RNaseH_epsilonretroviridae\|RNaseH;RNaseH_gammaretroviridae | HERVE−int;LTR/ERV1 | UBXN8;ENSP00000483433 | ≥50% | 4710 |
| Hsap38.chr9.32631428.32631838.− | Gag | gag;zf−CCHC_6 | . | TAF1L;ENSP00000418379 | | |
| Hsap38.chr12.8224015.8224431.− | Gag | gag;zf−CCHC_6 | . | FAM90A1;ENSP00000445418 | ≥50% | 3274 |
| Hsap38.chr12.42444627.42445106.+ | Pol | pol;RNase_H\|RNaseH;RNaseH_betaretroviridae | HERVK22−int;LTR/ERVK | PPHLN1;ENSP00000338510 | ≥10% | 2306 |
| Hsap38.chr12.42445855.42446199.+ | Pol | pol;IN_DBD_C\|INT;INT_betaretroviridae | HERVK22−int;LTR/ERVK | PPHLN1;ENSP00000477681 | ≥50% | |
| Hsap38.chr14.100880715.100884845.− | Pro | pro;gag−asp_proteas\|AP;AP_v_clade\|GAG;GAG_v_clade\|RT;RT_17_6\|RT;RT_cer2_3\|RT;RT_crm\|RT;RT_del\|RT;RT_gypsy\|RT;RT_maggy\|RT;RT_pyggy\|RT;RT_v_clade | (GGTG)n;Simple_repeat | RTL1;ENSP00000435342 | | 752 |
| Hsap38.chr19.18995047.18995343.− | Pro | pro;G−patch | . | SUGP2;ENSP00000470915 | ≥50% | |
| Hsap38.chr19.53013986.53015521.+ | Env | env;TLV_coat\|ENV;ENV_D− | HERV4_I−int;LTR/ERV1 | ERVV−1;ENSP00000473153 | | 1821 |
| Hsap38.chr19.53049147.53050856.+ | Env | env;TLV_coat\|ENV;ENV_D− | MER66−int;LTR/ERV1 | ERVV−2;ENSP00000472919 | | 4506 |
| Hsap38.chr19.58311871.58312203.+ | Pol | pol;IN_DBD_C\|INT;INT_alpharetroviridae\|INT;I | HERVK3−int;LTR/ERVK | ERVK3−1;ENSP00000489088 | | 6243 |
| Hsap38.chr19.23574222.23574482.− | Pol | INT;INT_betaretroviridae | HERVK9−int;LTR/ERVK | ZNF675;ENSP00000473217 | | |
| Hsap38.chrX.71401441.71401740.+ | Gag | gag;zf−CCHC_6 | MER8;DNA/TcMar−Tigger | TAF1;ENSP00000406549 | ≥50% | |

#### Chimpanzee EVE-ORFs

| ID | Domain | HMM | Repbase | Gene |
|---|---|---|---|---|
| Ptro214.chr3.132592082.132592324.- | Gag | GAG;GAG_lentiviridae | . | CNBP;ENSPTRP00000026477 |
| Ptro214.chr5.103437732.103438058.- | Pol | INT;GIN1 | . | GIN1;ENSPTRP00000054536 |
| Ptro214.chr5.27254075.27254353.+ | Pol | INT;INT_epsilonretroviridae\|INT;INT_gammaretroviridae | HERVE-int;LTR/ERV1 | TMEM161B;ENSPTRP00000029207 |
| Ptro214.chr6.11189490.11191109.- | Env | ENV;ENV_deltaretroviridae\|ENV;ENV_D-type_betaretroviridae\|ENV;ENV_gammaretroviridae\|ENV;ENV_retroviridae | MER50-int;LTR/ERV1 | ERVFRD-1;ENSPTRP00000058789 |
| Ptro214.chr7.92995230.92996849.- | Env | env;TLV_coat\|ENV;ENV_D-type betaretroviridae\| ENV; ENV_gammaretroviridae | HERV17-int;LTR/ERV1 | ERVW-1 |
| Ptro214.chr7.136045679.136046137.+ | Env | env;TLV_coat\|ENV;ENV_D-type_betaretroviridae\|ENV;ENV_gammaretroviridae | HERVI-int;LTR/ERV1 | AKR1B15;ENSPTRP00000058399 |
| Ptro214.chr7.95191982.95193214.+ | Gag | GAG;GAG_v_clade | (CCT)n;Simple_repeat\|(CAA)n;Simple_repeat | PEG10;ENSPTRP00000045828 |
| Ptro214.chr7.95193157.95194362.+ | Pro;G-patch | AP;AP_v_clade | (ACC)n;Simple_repeat\|(CCG)n;Simple_repeat | PEG10;ENSPTRP00000045828 |
| Ptro214.chr8.51811074.51811343.+ | Pol | RT;RT_alpharetroviridae\|RT;RT_betaretroviridae | AluYGibC1;SINE/Alu\|HERVK9-int;LTR/ERVK | RPL9;ENSPTRP00000061373 |
| Ptro214.chr12.8478239.8478652.- | Gag | gag;zf-CCHC_6 | . | FAM90A1;ENSPTRP00000050473 |
| Ptro214.chr19.19312408.19312704.- | Pro | pro;G-patch | . | SUGP2;ENSPTRP00000045722 |

#### Gorilla EVE-ORFs

| ID | Domain | HMM | Repbase | Gene |
|---|---|---|---|---|
| Ggor31.chr1.197650039.197650293.- | Pro | pro;G-patch | . | GPATCH2;ENSGGOP00000025170 |
| Ggor31.chr2a.71259560.71260618.- | Pro | AP;AP_DTG_ILG_template\|AP;AP_saspase | . | ASPRV1;ENSGGOP00000024375 |
| Ggor31.chr3.129088032.129088274.- | Gag | GAG;GAG_lentiviridae | . | CNBP;ENSGGOP00000009839 |
| Ggor31.chr5.85945229.85945555.- | Pol | INT;GIN1 | . | GIN1;ENSGGOP00000026207 |
| Ggor31.chr6.11566308.11567927.- | Env | ENV;ENV_deltaretroviridae\|ENV;ENV_D-type_betaretroviridae\|ENV;ENV_retroviridae | MER50-int;LTR/ERV1 | ERVFRD-1;ENSGGOP00000021598 |
| Ggor31.chr7.89729672.89731291.- | Env | env;TLV_coat\|ENV;ENV_D-type betaretroviridae\| ENV; ENV_gammaretroviridae | HERV17-int;LTR/ERV1 | ERVW-1 |
| Ggor31.chr7.103145635.103146048.- | Pol | RNaseH;RNaseH_betaretroviridae | (CGG)n;Simple_repeat\|HERVK9-int;LTR/ERVK | YBX1;ENSGGOP00000017832 |
| Ggor31.chr12.40175362.40175706.+ | Pol | pol;IN_DBD_C\|INT;INT_betaretroviridae | (GCAGCCCCC)n;Simple_repeat\|HERVK22-int;LTR/ERVK | PPHLN1;ENSGGOP00000017687 |
| Ggor31.chr19.19372895.19373191.- | Pro | pro;G-patch | . | SUGP2;ENSGGOP00000021642 |
| Ggor31.chrX.68663891.68664187.+ | Gag | gag;zf-CCHC_6 | MER8;DNA/TcMar-Tigger | TAF1;ENSGGOP00000015303 |

#### Orangtan EVE-ORFs

| ID | Domain | HMM | Repbase | Gene |
|---|---|---|---|---|
| Pabe2.chr1.32354186.32354437.+ | Pro | pro;G-patch | . | GPATCH2;ENSPPYG00000000216 |
| Pabe2.chr2a.40839531.40840589.+ | Pro | AP;AP_DTG_ILG_template\|AP;AP_saspase | . | ASPRV1;ENSPPYG00000012330 |
| Pabe2.chr3.108279426.108280022.- | Pol | INT;INT_alpharetroviridae\|INT;INT_betaretroviridae\|INT;INT_lentiviridae | HERVK9-int;LTR/ERVK | XYLB;ENSPPYG00000014007 |
| Pabe2.chr3.7275585.7275947.- | LINE | pro;dUTPase | L1MB1;LINE/L1 | NA;ENSPPYG00000013463 |
| Pabe2.chr5.103510614.103510940.- | Pol | INT;GIN1 | . | GIN1;ENSPPYG00000015666 |
| Pabe2.chr6.11476609.11478228.- | Env | ENV;ENV_deltaretroviridae\|ENV;ENV_D-type_betaretroviridae\|ENV;ENV_retroviridae | MER50-int;LTR/ERV1 | ERVFRD-1;ENSPPYG00000016231 |
| Pabe2.chr7.81421560.81422765.- | Pro | AP;AP_v_clade | (GGTGGC)n;Simple_repeat\|(GTG)n;Simple_repeat | NA;ENSPPYG00000029380 |
| Pabe2.chr7.83678858.83680477.+ | Env | env;TLV_coat\|ENV;ENV_D-type betaretroviridae\| ENV; ENV_gammaretroviridae | HERV17-int;LTR/ERV1 | ERVW-1 |
| Pabe2.chr8.7379820.7380095.- | Gag | gag;zf-CCHC_6 | . | NA;ENSPPYG00000018331 |
| Pabe2.chr9.29027142.29027552.+ | Gag | gag;zf-CCHC_6 | . | TAF1L;ENSPPYG00000019145 |
| Pabe2.chr11.61285274.61285540.+ | Pro | AP;AP_nix1 | . | NRIP3;ENSPPYG00000003499 |
| Pabe2.chr12.41813010.41813336.+ | Pol | pol;IN_DBD_C | HERVK22-int;LTR/ERVK | PPHLN1;ENSPPYG00000004423 |
| Pabe2.chr14.102426095.102430228.- | Pro:Gag | pro;gag-asp_proteas\|AP;AP_v_clade\|GAG;GAG_v_clade\|RT;RT_17_6\|RT;RT_cer2_3\|RT;RT_crm\|RT;RT_del\|RT;RT_gypsy\|RT;RT_maggy\|RT;RT_pygy\|RT;RT_v_clade | (GGTG)n;Simple_repeat | RTL1;ENSPPYG00000006149 |
| Pabe2.chr17.20152211.20152498.- | Pol | pol;RNase_H | . | NA;ENSPPYG00000008067 |
| Pabe2.chr19.54996417.54996668.- | Gag | gag;zf-CCHC_6 | . | NA;ENSPPYG00000010352 |
| Pabe2.chr19.60435534.60435866.+ | Pol | pol;IN_DBD_C\|INT;INT_alpharetroviridae\|INT;INT_betaretroviridae | HERVK3-int;LTR/ERVK | ERVK3-1;ENSPPYG00000029674 |
| Pabe2.chr21.44310326.44310838.- | Env | (env) | . | NA;ENSPPYG00000011460 |
| Pabe2.chr21.44310894.44311181.- | Pol | INT;INT_gammaretroviridae | HERVH48-int;LTR/ERV1 | NA;ENSPPYG00000011460 |
| Pabe2.chrX.68901666.68901965.+ | Gag | gag;zf-CCHC_6 | MER8;DNA/TcMar-Tigger | TAF1;ENSPPYG00000020453 |

#### Baboon EVE-ORFs

<table>
<tr><th>ID</th><th>Domain</th><th>HMM</th><th>Repbase</th><th>Gene</th></tr>
<tr><td>Panu2.chr3.107351109.107352314.–</td><td>Pro</td><td>AP;AP_v_clade</td><td>(GGC)n;Simple_repeat|(GTG)n;Simple_repeat</td><td>PEG10;ENSPANP00000010940</td></tr>
<tr><td>Panu2.chr3.107352257.107353555.–</td><td>Gag</td><td>GAG;GAG_v_clade</td><td>(GTT)n;Simple_repeat</td><td>PEG10;ENSPANP00000010940</td></tr>
<tr><td>Panu2.chr4.11017359.11018975.–</td><td>Env</td><td>ENV;ENV_gammaretroviridae|ENV;ENV_retroviridae</td><td>MER50–int;LTR/ERV1</td><td>ERVFRD–1;ENSPANP00000010950</td></tr>
<tr><td>Panu2.chr6.96802166.96802492.–</td><td>Pol</td><td>INT;GIN1</td><td>.</td><td>GIN1;ENSPANP00000011998</td></tr>
<tr><td>Panu2.chr7.157054388.157058521.–</td><td>Pro:Gag</td><td>pro;gag–asp_proteas|AP;AP_v_clade|GAG;GAG_v_clade|RT;RT_17_6|RT;RT_cer2_3|RT;RT_crm|RT;RT_del|RT;RT_gypsy|RT;RT_maggy|RT;RT_pyggy|RT;RT_pyret|RT;RT_v_clade</td><td>.</td><td>RTL1;ENSPANP00000000445</td></tr>
<tr><td>Panu2.chr11.83922569.83922811.+</td><td>Gag</td><td>GAG;GAG_lentiviridae</td><td>.</td><td>CNBP;ENSPANP00000003453</td></tr>
<tr><td>Panu2.chr12.36760606.36761160.+</td><td>Pro</td><td>pro;dUTPase|DUT;DUT_caulimoviruses|DUT;DUT_lentiviridae</td><td>.</td><td>NA;ENSPANP00000009281</td></tr>
<tr><td>Panu2.chr13.68869338.68870462.–</td><td>Pro</td><td>AP;AP_DTG_ILG_template|AP;AP_saspase</td><td>.</td><td>ASPRV1;ENSPANP00000007998</td></tr>
<tr><td>Panu2.chr14.1438165.1438500.–</td><td>Pro</td><td>pro;Asp|AP;AP_pepsins_A1a</td><td>.</td><td>NA;ENSPANP00000007098</td></tr>
<tr><td>Panu2.chr19.17555770.17556051.–</td><td>Pro</td><td>pro;G–patch</td><td>.</td><td>SUGP2;ENSPANP00000000126</td></tr>
<tr><td>Panu2.chrX.62805510.62805842.+</td><td>Gag</td><td>gag;zf–CCHC_6</td><td>MER8;DNA/TcMar–Tigger</td><td>TAF1;ENSPANP00000008327</td></tr>
</table>

#### Rhesus magaque EVE-ORFs

<table>
<tr><th>ID</th><th>Domain</th><th>HMM</th><th>Repbase</th><th>Gene</th></tr>
<tr><td>Mmul1.chr11.85671618.85671860.+</td><td>Gag</td><td>GAG;GAG_lentiviridae</td><td>.</td><td>CNBP;ENSMMUP00000015215</td></tr>
<tr><td>Mmul1.chr2.8788584.8789783.+</td><td>LINE</td><td>pol;RVT_1</td><td>L1_RS1;LINE/L1</td><td>NA;ENSMMUP00000040168</td></tr>
<tr><td>Mmul1.chr3.60406566.60408392.+</td><td>Env</td><td>ENV;ENV_gammaretroviridae</td><td>HERV3–int;LTR/ERV1</td><td>ERV3–1;ENSMMUP00000035333</td></tr>
<tr><td>Mmul1.chr4.110696699.11071315.–</td><td>Env</td><td>ENV;ENV_deltaretroviridae|ENV;ENV_D–type_betaretroviridae|ENV;ENV_retroviridae</td><td>MER50–int;LTR/ERV1</td><td>ERVFRD–1</td></tr>
<tr><td>Mmul1.chr5.128640286.128640867.+</td><td>LINE</td><td>pol;RVT_1</td><td>L1–2_Cja;LINE/L1</td><td>NA;ENSMMUP00000030240</td></tr>
<tr><td>Mmul1.chr6.154841863.154843983.+</td><td>LINE</td><td>pol;RVT_1</td><td>L1_RS1;LINE/L1</td><td>NA;ENSMMUP00000033787</td></tr>
<tr><td>Mmul1.chr6.99336613.99336939.–</td><td>Pol</td><td>INT;GIN1</td><td>.</td><td>GIN1;ENSMMUP00000010230</td></tr>
<tr><td>Mmul1.chr7.164164659.164168792.–</td><td>Pro:Gag</td><td>pro;gag–asp_proteas|AP;AP_v_clade|GAG;GAG_v_clade|RT;RT_17_6|RT;RT_cer2_3|RT;RT_crm|RT;RT_del|RT;RT_gypsy|RT;RT_maggy|RT;RT_pyggy|RT;RT_pyret|RT;RT_v_clade</td><td>.</td><td>RTL1;ENSMMUP00000007234</td></tr>
<tr><td>Mmul1.chr13.70221349.70222476.–</td><td>Pro</td><td>AP;AP_DTG_ILG_template|AP;AP_saspase</td><td>.</td><td>ASPRV1;ENSMMUP00000011804</td></tr>
<tr><td>Mmul1.chr14.1669280.1669618.–</td><td>Pro</td><td>pro;Asp|AP;AP_pepsins_A1a</td><td>.</td><td>NA;ENSMMUP00000005169</td></tr>
<tr><td>Mmul1.chr19.59382968.59383237.+</td><td>Gag</td><td>gag;zf–CCHC_6</td><td>.</td><td>NA;ENSMMUP00000021136</td></tr>
<tr><td>Mmul1.chr19.64064515.64065216.+</td><td>Pol</td><td>INT;INT_betaretroviridae</td><td>AluYRb3;SINE/Alu|HERVK3–int;LTR/ERVK</td><td>ERVK3–1;ENSMMUP00000027005</td></tr>
<tr><td>Mmul1.chrX.123292284.123293480.+</td><td>LINE</td><td>pol;RVT_1</td><td>L1_RS1;LINE/L1</td><td>NA;ENSMMUP00000040834</td></tr>
<tr><td>Mmul1.chrX.70302480.70302812.+</td><td>Gag</td><td>gag;zf–CCHC_6</td><td>MER8;DNA/TcMar–Tigger</td><td>TAF1;ENSMMUP00000007484</td></tr>
</table>

#### Marmoset EVE-ORFs

<table>
<tr><th>ID</th><th>Domain</th><th>HMM</th><th>Repbase</th><th>Gene</th></tr>
<tr><td>Cjac321.chr4.17531888.17533507.+</td><td>Env</td><td>ENV;ENV_D-type betaretrovidae |ENV;ENV_retroviridae</td><td>MER50–int;LTR/ERV1</td><td>ERVFRD-1:ENSCJAP00000040973</td></tr>
<tr><td>Cjac321.chr2.104319490.104319867.+</td><td>Pol</td><td>INT;GIN1</td><td>.</td><td>GIN1;ENSCJAP00000016966</td></tr>
<tr><td>Cjac321.chr6.70290164.70292944.–</td><td>LINE</td><td>pol;RVT_1</td><td>L1–1_Cja;LINE/L1|Sat–1_TSy;Satellite</td><td>LYPD6B;ENSCJAP00000005582</td></tr>
<tr><td>Cjac321.chr13.6531633.6531890.+</td><td>Gag</td><td>gag;zf–CCHC_6</td><td>.</td><td>NA;ENSCJAP00000007145</td></tr>
<tr><td>Cjac321.chr15.67817302.67817544.+</td><td>Gag</td><td>GAG;GAG_lentiviridae</td><td>.</td><td>CNBP;ENSCJAP00000032138</td></tr>
<tr><td>Cjac321.chr22.17949261.17949596.–</td><td>Pro</td><td>pro;G–patch</td><td>.</td><td>SUGP2;ENSCJAP00000036644</td></tr>
<tr><td>Cjac321.chrX.63332816.63333115.+</td><td>Gag</td><td>gag;zf–CCHC_6</td><td>MER8;DNA/TcMar–Tigger</td><td>NA;ENSCJAP00000043164</td></tr>
</table>

### Mouse EVE-ORFs

<table>
<tr><th>ID</th><th>Domain</th><th>HMM</th><th>Repbase</th><th>Gene</th><th>Status</th><th>Dist_TSS</th></tr>
<tr><td>Mmus38.chr1.131672160.131672417.+</td><td>Pro</td><td>pro;Asp|AP;AP_pepsins_A1a</td><td>.</td><td>Ctse;ENSMUSP00000108030</td><td></td><td>0</td></tr>
<tr><td>Mmus38.chr1.173568891.173569334.-</td><td>Pol</td><td>INT;INT_betaretroviridae</td><td>ERVB2_1-I_MM;LTR/ERVK</td><td>Pydc4;ENSMUSP00000117222</td><td></td><td></td></tr>
<tr><td>Mmus38.chr1.173695398.173695781.+</td><td>Pol</td><td>INT;INT_betaretroviridae</td><td>ERVB2_1-I_MM;LTR/ERVK</td><td>Pydc3;ENSMUSP00000128958</td><td></td><td></td></tr>
<tr><td>Mmus38.chr1.78497642.78498025.-</td><td>Env</td><td>env;TLV_coat|ENV;ENV_D-type_betaretroviridae|ENV;ENV_gammaretroviridae</td><td>MMERGLN-int;LTR/ERV1</td><td>BC035947;ENSMUSP00000132488</td><td></td><td></td></tr>
<tr><td>Mmus38.chr1.78497809.78500529.-</td><td>Pol</td><td>pol;rve|env;TLV_coat|ENV;ENV_deltaretroviridae|ENV;ENV_D-type_betaretroviridae|ENV;ENV_gammaretroviridae|ENV;ENV_retroviridae|INT;INT_athila|INT;INT_b_clade|INT;INT_csrn1|INT;INT_deltaretroviridae|INT;INT_epsilonretroviridae|INT;INT_gammaretroviridae|INT;INT_gmr1|INT;INT_lentiviridae|INT;INT_osvaldo|INT;INT_spumaretroviridae|INT;INT_tat</td><td>MMERGLN-int;LTR/ERV1</td><td>BC035947;ENSMUSP00000132488</td><td></td><td></td></tr>
<tr><td>Mmus38.chr1.97792275.97792568.+</td><td>Pol</td><td>INT;GIN1</td><td>.</td><td>Gin1;ENSMUSP00000027571</td><td></td><td></td></tr>
<tr><td>Mmus38.chr2.155021472.155024216.+</td><td>Pol</td><td>pol;IN_DBD_C|pol;Integrase_Zn|pol;RNase_H|pol;rve|pol;RVT_1|pol;RVT_thumb|INT;INT_alpharetroviridae|INT;INT_b_clade|INT;INT_betaretroviridae|INT;INT_deltaretroviridae|INT;INT_gammaretroviridae|INT;INT_lentiviridae|INT;INT_spumaretroviridae|RNaseH;RNaseH_betaretroviridae|RT;RT_alpharetroviridae|RT;RT_betaretroviridae|RT;RT_del|RT;RT_deltaretroviridae|RT;RT_epsilonretroviridae|RT;RT_lentiviridae|RT;RT_spumaretroviridae</td><td>ERVB4_1-I_MM;LTR/ERVK</td><td>Gm14226;ENSMUSP00000122157</td><td></td><td>2603</td></tr>
<tr><td>Mmus38.chr2.155024056.155026182.+</td><td>Env</td><td>env;TLV_coat|ENV;ENV_deltaretroviridae|ENV;ENV_D-type_betaretroviridae|ENV;ENV_gammaretroviridae|ENV;ENV_retroviridae</td><td>ERVB4_1-I_MM;LTR/ERVK</td><td>Gm14226;ENSMUSP00000122157</td><td></td><td>4968</td></tr>
<tr><td>Mmus38.chr3.137672404.137675022.+</td><td>LINE</td><td>pol;RVT_1</td><td>L1Md_A;LINE/L1</td><td>Gm21962;ENSMUSP00000126626</td><td></td><td></td></tr>
<tr><td>Mmus38.chr4.150904482.150905285.-</td><td>Pro</td><td>pro;dUTPase</td><td>MERVL-int;LTR/ERVL</td><td>Park7;ENSMUSP00000122265</td><td></td><td>2215</td></tr>
<tr><td>Mmus38.chr5.134558243.134560099.-</td><td>Env</td><td>ENV;ENV_gammaretroviridae|ENV;ENV_retroviridae</td><td>RodERV21-int;LTR/ERV1</td><td>Syna;ENSMUSP00000116437</td><td>SINGL</td><td>3742</td></tr>
<tr><td>Mmus38.chr5.23701149.23703218.-</td><td>Env</td><td>env;TLV_coat|ENV;ENV_deltaretroviridae|ENV;ENV_D-type_betaretroviridae|ENV;ENV_gammaretroviridae|ENV;ENV_retroviridae</td><td>MuLV-int;KTR/ERV1</td><td>Fv4;NA</td><td>SINGL</td><td>0</td></tr>
<tr><td>Mmus38.chr6.129859271.129863149.+</td><td>LINE</td><td>pol;RVT_1</td><td>L1Md_T;LINE/L1|L1Md_Gf;LINE/L1</td><td>Gm17631;ENSMUSP00000130863</td><td></td><td></td></tr>
<tr><td>Mmus38.chr6.4754122.4755348.+</td><td>Gag</td><td>GAG;GAG_v_clade</td><td>(CAA)n;Simple_repeat|(GAT)n;Simple_repeat</td><td>Peg10;ENSMUSP00000127306</td><td></td><td>0</td></tr>
<tr><td>Mmus38.chr6.4755117.4757237.+</td><td>Pro</td><td>AP;AP_v_clade</td><td>(ATCTGC)n;Simple_repeat|(CCA)n;Simple_repeat</td><td>Peg10;ENSMUSP00000127306</td><td></td><td>0</td></tr>
<tr><td>Mmus38.chr6.86628075.86629190.+</td><td>Pro</td><td>AP;AP_DTG_ILG_template|AP;AP_saspase</td><td>.</td><td>Asprv1;ENSMUSP00000046121</td><td>SINGL</td><td>0</td></tr>
<tr><td>Mmus38.chr6.87845121.87845363.-</td><td>Gag</td><td>GAG;GAG_lentiviridae</td><td>.</td><td>Cnbp;ENSMUSP00000109247</td><td></td><td></td></tr>
<tr><td>Mmus38.chr7.104271650.104273401.-</td><td>Gag</td><td>gag;Gag_p24|gag;zf-CCHC_5|GAG;GAG_betaretroviridae|GAG;GAG_lentiviridae</td><td>IAPEz-int;LTR/ERVK</td><td>Trim5;ENSMUSP00000050084</td><td></td><td></td></tr>
<tr><td>Mmus38.chr7.142380732.142380977.-</td><td>Pro</td><td>pro;Asp|AP;AP_pepsins_A1a</td><td>.</td><td>Ctsd;ENSMUSP00000063904</td><td></td><td>0</td></tr>
<tr><td>Mmus38.chr7.143635667.143636467.-</td><td>Pol</td><td>pol;RNase_H</td><td>MERVL-int;LTR/ERVL</td><td>Tnfrsf22;ENSMUSP00000126384</td><td></td><td></td></tr>
<tr><td>Mmus38.chr8.70259617.70259880.+</td><td>Pro</td><td>pro;G-patch</td><td>.</td><td>Sugp2;ENSMUSP00000128029</td><td></td><td></td></tr>
<tr><td>Mmus38.chr12.109590172.109595502.-</td><td>Pro:Gag</td><td>pro;gag-asp_proteas|AP;AP_v_clade|GAG;GAG_v_clade|RT;RT_17_6|RT;RT_crm|RT;RT_del|RT;RT_pyret|RT;RT_reina|RT;RT_v_clade</td><td>(TCC)n;Simple_repeat|(CTCAGC)n;Simple_repeat</td><td>Rtl1;ENSMUSP00000115957</td><td>SINGL</td><td>0</td></tr>
<tr><td>Mmus38.chr14.43930744.43931031.+</td><td>Pol</td><td>pol;IN_DBD_C|INT;INT_betaretroviridae</td><td>IAPEY4_I-int;LTR/ERVK</td><td>Gm8113;ENSMUSP00000125038</td><td></td><td>5402</td></tr>
<tr><td>Mmus38.chr14.69290950.69292893.-</td><td>Env</td><td>ENV;ENV_retroviridae</td><td>RodERV21-int;LTR/ERV1</td><td>Synb;ENSMUSP00000061107</td><td>SINGL</td><td>4431</td></tr>
<tr><td>Mmus38.chr14.73660751.73663234.+</td><td>LINE</td><td>pol;RVT_1</td><td>L1Md_A;LINE/L1</td><td>Gm21750;ENSMUSP00000096471</td><td></td><td></td></tr>
<tr><td>Mmus38.chr16.15317458.15321357.+</td><td>LINE</td><td>pol;RVT_1</td><td>L1Md_T;LINE/L1</td><td>Gm21897;ENSMUSP00000135960</td><td></td><td></td></tr>
<tr><td>Mmus38.chrX.101563477.101563740.+</td><td>Gag</td><td>gag;zf-CCHC_6</td><td>.</td><td>Taf1;ENSMUSP00000098895</td><td></td><td></td></tr>
<tr><td>Mmus38.chrX.94356887.94357219.-</td><td>Gag</td><td>gag;zf-CCHC_6</td><td>(CT)n;Simple_repeat</td><td>Fam90a1b;ENSMUSP00000109536</td><td></td><td></td></tr>
<tr><td>Mmus38.chrY.70293570.70293914.-</td><td>Pol</td><td>pol;rve|INT;INT_epsilonretroviridae|INT;INT_gammaretroviridae</td><td>MuRRS4-int;LTR/ERV1</td><td>Gm29423;ENSMUSP00000140691</td><td></td><td></td></tr>
</table>

#### Rat EVE-ORFs

<table>
<tr><th>ID</th><th>Domain[</th><th>HMM</th><th>Repbase</th><th>Gene</th></tr>
<tr><td>Rnor50.chr1.222441769.222442107.−</td><td>Pro</td><td>pro;Asp|AP;AP_pepsins_A1a</td><td>.</td><td>Ctsd;ENSRNOP00000027407</td></tr>
<tr><td>Rnor50.chr1.51190516.51190869.+</td><td>Pro</td><td>pro;dUTPase|DUT;DUT_caulimoviruses|DUT;DUT_lentiviridae</td><td>.</td><td>AABR06002285.1;ENSRNOP00000023387</td></tr>
<tr><td>Rnor50.chr1.73779954.73782599.−</td><td>LINE</td><td>pol;RVT_1</td><td>L1_Rat2;LINE/L1</td><td>AABR06003466.1;ENSRNOP00000012225</td></tr>
<tr><td>Rnor50.chr2.132087070.132091020.−</td><td>LINE</td><td>pol;RVT_1</td><td>RNHAL1;LINE/L1</td><td>AABR06015425.1;ENSRNOP00000067570</td></tr>
<tr><td>Rnor50.chr2.221363325.221367224.+</td><td>LINE</td><td>pol;RVT_1</td><td>L1_Rn;LINE/L1</td><td>AABR06019553.1;ENSRNOP00000064593</td></tr>
<tr><td>Rnor50.chr4.185001418.185001660.−</td><td>Gag</td><td>GAG;GAG_lentiviridae</td><td>.</td><td>Cnbp;ENSRNOP00000013884</td></tr>
<tr><td>Rnor50.chr4.221377781.221378269.+</td><td>LINE</td><td>pol;RVT_1</td><td>L1_Rn;LINE/L1</td><td>Mug2;ENSRNOP00000064788</td></tr>
<tr><td>Rnor50.chr6.137174293.137175420.+</td><td>LINE</td><td>pol;RVT_1</td><td>L1_Rat2;LINE/L1</td><td>LOC500712;ENSRNOP00000029694</td></tr>
<tr><td>Rnor50.chr6.142874258.142877095.−</td><td>Gag</td><td>GAG;GAG_v_clade|RT;RT_crm|RT;RT_del|RT;RT_pyret|RT;RT_reina|RT;RT_v_clade</td><td>(TCCTCA)n;Simple_repeat|(ATCTTC)n;Simple_repeat</td><td>Rtl1;ENSRNOP00000066132</td></tr>
<tr><td>Rnor50.chr7.95855614.95857533.+</td><td>Pol</td><td>pol;RVT_1</td><td>L1_Rn;LINE/L1</td><td>Col14a1;ENSRNOP00000063778</td></tr>
<tr><td>Rnor50.chr9.110750764.110751060.−</td><td>Pol</td><td>INT;GIN1</td><td>.</td><td>Gin1;ENSRNOP00000016028</td></tr>
<tr><td>Rnor50.chr9.52775737.52777266.−</td><td>Pol</td><td>pol;IN_DBD_C|pol;Integrase_Zn|pol;RNase_H|pol;rve|INT;INT_alpharetroviridae|INT;INT_betaretroviridae|INT;INT_deltaretroviridae|INT;INT_gammaretroviridae|INT;INT_lentiviridae|INT;INT_spumaretroviridae|RNaseH;RNaseH_betaretroviridae</td><td>RNLTR4d_I;LTR/ERVK</td><td>Ormdl1;ENSRNOP00000061507</td></tr>
<tr><td>Rnor50.chr9.52779885.52780949.−</td><td>Gag</td><td>GAG;GAG_betaretroviridae</td><td>RNLTR4d_I;LTR/ERVK</td><td>Ormdl1;ENSRNOP00000061507</td></tr>
<tr><td>Rnor50.chr10.70117964.70118332.−</td><td>Gag</td><td>gag;Gag_p24|GAG;GAG_betaretroviridae</td><td>MYSERV_Rn-int;LTR/ERVK</td><td>Rn50_10_0702.5;ENSRNOP00000012874</td></tr>
<tr><td>Rnor50.chr12.27165685.27167544.+</td><td>Env</td><td>ENV;ENV_gammaretroviridae|ENV;ENV_retroviridae</td><td>RodERV21-int;LTR/ERV1</td><td>Syna;ENSRNOP00000067387</td></tr>
<tr><td>Rnor50.chr13.116482035.116484032.−</td><td>Pol</td><td>pol;RVT_1</td><td>L1_Rat1;LINE/L1</td><td>Hsd11b1;ENSRNOP00000050581</td></tr>
<tr><td>Rnor50.chr15.54904810.54906750.−</td><td>Env</td><td>ENV;ENV_gammaretroviridae|ENV;ENV_retroviridae</td><td>RodERV21-int;LTR/ERV1</td><td>Synb;ENSRNOG00000021823</td></tr>
<tr><td>Rnor50.chr16.20764085.20764363.−</td><td>Pro</td><td>pro;G-patch</td><td>.</td><td>Sugp2;ENSRNOP00000027457</td></tr>
<tr><td>Rnor50.chr17.46488518.46491379.+</td><td>Pol</td><td>pol;RVT_1</td><td>RNHAL1;LINE/L1|L1_Rn;LINE/L1</td><td>AABR06091708.2;ENSRNOP00000066488</td></tr>
<tr><td>Rnor50.chr17.47415263.47417911.+</td><td>Pol</td><td>pol;RVT_1</td><td>RNHAL1;LINE/L1</td><td>Pou6f2;ENSRNOP00000017713</td></tr>
<tr><td>Rnor50.chr19.32976355.32980209.−</td><td>LINE</td><td>pol;RVT_1</td><td>L1_Rn;LINE/L1</td><td>AABR06097607.1;ENSRNOP00000065391</td></tr>
<tr><td>Rnor50.chr19.47076640.47077344.+</td><td>Pol</td><td>pol;rve|INT;INT_betaretroviridae|INT;INT_deltaretroviridae|INT;INT_gammaretroviridae</td><td>RMER16-int;LTR/ERVK</td><td>AABR06098457.1;ENSRNOP00000038976</td></tr>
</table>

### Rabbit EVE-ORFs

<table>
<tr><th>ID</th><th>Domain</th><th>HMM</th><th>Repbase</th><th>Gene</th></tr>
<tr><td>Ocun2.chr1.182294471.182297680.-</td><td>Pol</td><td>pol;RVT_1</td><td>Sat-1_TSy;Satellite|L1A2_OC;LINE/L1</td><td>NA;ENSOCUG00000029631</td></tr>
<tr><td>Ocun2.chr1.192706896.192709235.+</td><td>LINE</td><td>pol;RVT_1</td><td>L1A_Oc;LINE/L1|Sat-1_TSy;Satellite|L1A2_OC;LINE/L1</td><td>NA;ENSOCUG00000029694</td></tr>
<tr><td>Ocun2.chr2.116246603.116247574.+</td><td>Pro</td><td>AP;AP_DTG_ILG_template|AP;AP_saspase</td><td>.</td><td>ASPRV1;ENSOCUG00000015830</td></tr>
<tr><td>Ocun2.chr21.12630468.12634019.-</td><td>Pol</td><td>pol;IN_DBD_C|pol;rve|INT;INT_alpharetroviridae|INT;INT_betaretroviridae|INT;INT_deltaretroviridae|INT;INT_gammaretroviridae|INT;INT_lentiviridae|INT;INT_spumaretroviridae|RNaseH;RNaseH_betaretroviridae</td><td>ERV2-2_OC-I;LTR/ERVK</td><td>NA;ENSOCUG00000029571</td></tr>
<tr><td>Ocun2.chr7.158233001.158236747.+</td><td>LINE</td><td>pol;RVT_1</td><td>L1A_Oc;LINE/L1|Sat-1_TSy;Satellite</td><td>NA;ENSOCUG00000029715</td></tr>
<tr><td>Ocun2.chr7.77715477.77718938.-</td><td>LINE</td><td>pol;RVT_1</td><td>Sat-1_TSy;Satellite|L1A_Oc;LINE/L1</td><td>NA;ENSOCUG00000029595</td></tr>
<tr><td>Ocun2.chr9.7824836.7825078.+</td><td>Gag</td><td>GAG;GAG_lentiviridae</td><td>.</td><td>CNBP;ENSOCUG00000008696</td></tr>
<tr><td>Ocun2.chr10.34278491.34279759.-</td><td>Pro</td><td>AP;AP_v_clade</td><td>.</td><td>PEG10;ENSOCUG00000027535</td></tr>
<tr><td>Ocun2.chr10.34279045.34279962.-</td><td>Gag</td><td>GAG;GAG_v_clade</td><td>.</td><td>PEG10;ENSOCUG00000027535</td></tr>
<tr><td>Ocun2.chr11.22459697.22460035.-</td><td>Pol</td><td>INT;GIN1</td><td>.</td><td>GIN1;ENSOCUG00000025993</td></tr>
<tr><td>Ocun2.chr12.5569750.5572134.-</td><td>Pol</td><td>pol;IN_DBD_C|pol;Integrase_Zn|pol;RNase_H|pol;rve|pol;RVT_1|pol;RVT_thumb|INT;INT_a_clade|INT;INT_alpharetroviridae|INT;INT_b_clade|INT;INT_betaretroviridae|INT;INT_deltaretroviridae|INT;INT_gammaretroviridae|INT;INT_lentiviridae|INT;INT_spumaretroviridae|RNaseH;RNaseH_betaretroviridae|RT;RT_alpharetroviridae|RT;RT_betaretroviridae|RT;RT_deltaretroviridae|RT;RT_gammaretroviridae|RT;RT_lentiviridae</td><td>ERVH_OC_I-int;LTR/ERVK</td><td>NA;ENSOCUG00000029713</td></tr>
<tr><td>Ocun2.chr12.5571980.5572339.-</td><td>Pol</td><td>RT;RT_alpharetroviridae|RT;RT_betaretroviridae|RT;RT_epsilonretroviridae|RT;RT_gammaretroviridae|RT;RT_lentiviridae</td><td>ERVH_OC_I-int;LTR/ERVK</td><td>NA;ENSOCUG00000029713</td></tr>
<tr><td>Ocun2.chr12.99949107.99950864.+</td><td>Env</td><td>env;TLV_coat|ENV;ENV_deltaretroviridae|ENV;ENV_D-type_betaretroviridae|ENV;ENV_gammaretroviridae|ENV;ENV_retroviridae</td><td>MacERV4_int-int;LTR/ERVK|KORV_I-int;LTR/ERV1</td><td>ORY1;ENSOCUG00000029640</td></tr>
<tr><td>Ocun2.chr13.102787390.102790377.-</td><td>LINE</td><td>pol;RVT_1</td><td>Sat-1_TSy;Satellite|L1A_Oc;LINE/L1</td><td>NA;ENSOCUG00000029537</td></tr>
<tr><td>Ocun2.chr13.24336471.24339437.-</td><td>LINE</td><td>pol;RVT_1</td><td>Sat-1_TSy;Satellite|L1A_Oc;LINE/L1</td><td>NA;ENSOCUG00000029477</td></tr>
<tr><td>Ocun2.chr18.22458305.22462174.-</td><td>LINE</td><td>pol;RVT_1</td><td>L1C_OC;LINE/L1|Sat-1_TSy;Satellite|L1A_Oc;LINE/L1</td><td>NA;ENSOCUG00000029155</td></tr>
<tr><td>Ocun2.chrX.16160870.16163458.+</td><td>Pol</td><td>pol;IN_DBD_C|pol;Integrase_Zn|pol;RNase_H|pol;rve|pol;RVT_1|pol;RVT_thumb|INT;INT_a_clade|INT;INT_alpharetroviridae|INT;INT_betaretroviridae|INT;INT_deltaretroviridae|INT;INT_gammaretroviridae|INT;INT_lentiviridae|INT;INT_spumaretroviridae|RNaseH;RNaseH_betaretroviridae|RT;RT_17_6|RT;RT_alpharetroviridae|RT;RT_badnavirus|RT;RT_betaretroviridae|RT;RT_deltaretroviridae|RT;RT_epsilonretroviridae|RT;RT_gammaretroviridae|RT;RT_lentiviridae|RT;RT_spumaretroviridae</td><td>ERVH_OC_I-int;LTR/ERVK</td><td>NA;ENSOCUG00000029748</td></tr>
<tr><td>Ocun2.chrX.49877023.49877274.+</td><td>Gag</td><td>gag;zf-CCHC_6</td><td>.</td><td>NA;ENSOCUG00000015656</td></tr>
<tr><td>Ocun2.chrX.62570394.62572061.+</td><td>Env</td><td>env;TLV_coat|ENV;ENV_deltaretroviridae|ENV;ENV_D-type_betaretroviridae|ENV;ENV_retroviridae</td><td>MacERV4_int-int;LTR/ERVK|KORV_I-int;LTR/ERV1</td><td>NA;ENSOCUG00000029381</td></tr>
</table>

#### Dog EVE-ORFs

| ID | Domain | HMM | Repbase | Gene |
|---|---|---|---|---|
| Cfam31.chr2.10125398.10125826.- | Pol | INT;INT_epsilonretroviridae\|INT;INT_gammaretroviridae | CfERV1−int;LTR/ERV1 | NA;ENSCAFP00000037385 |
| Cfam31.chr2.10125562.10125852.- | Pol | RT;RT_epsilonretroviridae\|RT;RT_gammaretroviridae | CfERV1−int;LTR/ERV1 | NA;ENSCAFP00000037385 |
| Cfam31.chr2.10125762.10126721.- | Pro | AP;AP_gammaretroviridae\|AP;AP_retropepsins\|RT;RT_epsilonretroviridae\|RT;RT_gammaretroviridae | CfERV1−int;LTR/ERV1 | NA;ENSCAFP00000037385 |
| Cfam31.chr3.8115754.8116047.+ | Pol | INT;GIN1 | . | GIN1;ENSCAFP00000011199 |
| Cfam31.chr3.82194695.82195876.- | Env | env;TLV_coat\|ENV;ENV_D−type_betaretroviridae\|ENV;ENV_gammaretroviridae\|ENV;ENV_retroviridae | CfERVF1−int;LTR/ERV1 | NA;ENSCAFP00000039733 |
| Cfam31.chr3.82196099.82198096.- | Pol | pol;RNase_H\|pol;rve\|INT;INT_b_clade\|INT;INT_betaretroviridae\|INT;INT_csrn1\|INT;INT_deltaretroviridae\|INT;INT_epsilonretroviridae\|INT;INT_gammaretroviridae\|INT;INT_gmr1\|INT;INT_lentiviridae\|INT;INT_spumaretroviridae\|INT;INT_tat\|RNaseH;RNaseH_epsilonretroviridae\|RNaseH;RNaseH_gammaretroviridae\|RNaseH;RNaseH_spumaretroviridae | CfERVF1−int;LTR/ERV1 | NA;ENSCAFP00000039733 |
| Cfam31.chr3.83844409.83845833.+ | Env | ENV;ENV_retroviridae | CarERV3−int;LTR?ERV1 | CAR1;NA |
| Cfam31.chr5.963770.964429.+ | LINE | pol;RVT_1 | L1_Canis1;LINE/L1 | NA;ENSCAFP00000042660 |
| Cfam31.chr8.69103085.69106390.- | Pro | AP;AP_v_clade\|GAG;GAG_v_clade\|RT;RT_17_6\|RT;RT_crm\|RT;RT_del\|RT;RT_gypsy\|RT;RT_maggy\|RT;RT_pyggy\|RT;RT_pyret\|RT;RT_reina\|RT;RT_v_clade | (CTC)n;Simple_repeat | RTL1;ENSCAFP00000026463 |
| Cfam31.chr8.8320179.8322245.- | LINE | pol;RVT_1 | L1_Cf;LINE/L1 | NA;ENSCAFP00000033727 |
| Cfam31.chr10.3285576.3286484.+ | Env | env;TLV_coat\|ENV;ENV_D−type_betaretroviridae\|ENV;ENV_gammaretroviridae\|ENV;ENV_retroviridae | CfERV1−int;LTR/ERV1 | NA;ENSCAFP00000037516 |
| Cfam31.chr10.68586978.68587886.- | Pro | AP;AP_DTG_ILG_template\|AP;AP_saspase | (CCGTCC)n;Simple_repeat\|(GCAGGT)n;Simple_repeat | ASPRV1;ENSCAFP0000000492[illegible] |
| Cfam31.chr12.32255437.32256183.+ | Env | env;GP41\|ENV;ENV_B−type_betaretroviridae | (T)n;Simple_repeat | NA;ENSCAFP00000042124 |
| Cfam31.chr12.49629127.49631124.- | Env | env;GP41\|ENV;ENV_B−type_betaretroviridae\|ENV;ENV_retroviridae | . | NA;ENSCAFP00000039835 |
| Cfam31.chr14.20127798.20128877.+ | Gag | GAG;GAG_v_clade | (CCT)n;Simple_repeat | PEG10;ENSCAFP00000041426 |
| Cfam31.chr14.20128379.20129443.+ | Pro | AP;AP_v_clade | . | PEG10;ENSCAFP00000041426 |
| Cfam31.chr16.54089590.54089898.- | Gag | gag;zf−CCHC_6 | . | NA;ENSCAFP00000034730 |
| Cfam31.chr18.46013222.46013488.- | Pro | pro;Asp\|AP;AP_pepsins_A1a | . | NA;ENSCAFP00000014791 |
| Cfam31.chr20.2926592.2926834.+ | Gag | GAG;GAG_lentiviridae | . | CNBP;ENSCAFP00000006421 |
| Cfam31.chr20.44128813.44129130.+ | Pro | pro;G−patch | . | SUGP2;ENSCAFP00000021327 |
| Cfam31.chr26.28604208.28605758.+ | Gag | gag;Gag_p30 | MER66−int;LTR/ERV1\|(CCAGCC)n;Simple_repeat | NA;ENSCAFP00000037400 |
| Cfam31.chr35.24271063.24272484.+ | Env | env;TLV_coat\|ENV;ENV_gammaretroviridae\|ENV;ENV_retroviridae | MER66−int;LTR/ERV1 | NA;ENSCAFP00000038239 |
| Cfam31.chr35.24993046.24993933.- | Pol | pol;rve\|INT;INT_b_clade\|INT;INT_deltaretroviridae\|INT;INT_epsilonretroviridae\|INT;INT_gammaretroviridae\|INT;INT_gmr1\|INT;INT_lentiviridae\|INT;INT_osvaldo\|INT;INT_spumaretroviridae | CfERV1−int;LTR/ERV1 | NA;ENSCAFP00000042892 |
| Cfam31.chrX.55720071.55720400.+ | Gag | gag;zf−CCHC_6 | . | NA;ENSCAFP00000025204 |

#### Cat EVE-ORFs

| ID | Domain | HMM | Repbase | Gene |
|---|---|---|---|---|
| Fcat62.chrA1.163926192.163926512.- | Pol | Pol;RVT_1 | . | GIN1;ENSFCAP00000025585 |
| Fcat62.chrA2.98418392.98419471.+ | Gag | GAG;GAG_v_clade | (CCT)n;Simple_repeat | PEG10;ENSFCAP00000024585 |
| Fcat62.chrA2.13751269.13751583.- | Pro | pro;G−patch | . | SUGP2;ENSFCAP00000009055 |
| Fcat62.chrB1.138206424.138207278.- | LINE | pol;RVT_1 | L1−2_Fc;LINE/L1 | NA;ENSFCAP00000017244 |
| Fcat62.chrB1.183859927.183861351.+ | Env | ENV;ENV_gammaretroviridae | CarERV3−int;LTR?ERV1 | CAR1; ENSFCAG00000028607 |
| Fcat62.chrB2.3491740.3493038.- | Env | ENV;ENV_gammaretroviridae\|ENV;ENV_retroviridae | CarERV2−int;LTR/ERV1 | NA;ENSFCAP00000024081 |
| Fcat62.chrA3.89267109.89268092.− | Pro | AP;AP_DTG_ILG_template\|AP;AP_saspase | . | ASPRV1;ENSFCAP00000017297 |
| Fcat62.chrB4.47615806.47616552.+ | Gag | GAG;GAG_gammaretroviridae | ERV1−3_FCa−I;LTR/ERV1 | NA;ENSFCAP00000025368 |
| Fcat62.chrB4.47616233.47616805.+ | Gag | GAG;GAG_gammaretroviridae | ERV1−3_FCa−I;LTR/ERV1 | NA;ENSFCAP00000025368 |
| Fcat62.chrB4.47618106.47618432.+ | Gag | gag;Gag_p30\|GAG;GAG_gammaretroviridae | ERV1−3_FCa−I;LTR/ERV1 | NA;ENSFCAP00000025368 |
| Fcat62.chrB4.47618264.47618530.+ | Gag | gag;Gag_p30\|GAG;GAG_gammaretroviridae | ERV1−3_FCa−I;LTR/ERV1 | NA;ENSFCAP00000025368 |
| Fcat62.chrB4.47618436.47619767.+ | Gag | gag;Gag_p30\|pol;rve\|GAG;GAG_gammaretroviridae\|INT;INT_b_clade\|INT;INT_epsilonretroviridae\|INT;INT_gammaretroviridae\|INT;INT_gmr1\|INT;INT_spumaretroviridae | ERV1−3_FCa−I;LTR/ERV1 | NA;ENSFCAP00000025368 |
| Fcat62.chrB4.47619368.47620531.+ | Env | env;TLV_coat | ERV1−3_FCa−I;LTR/ERV1 | NA;ENSFCAP00000025368 |
| Fcat62.chrC1.93051406.93051657.+ | Pro | AP;AP_pepsins_A1a | . | NA;ENSFCAP00000019684 |
| Fcat62.chrD1.104121318.104122013.- | Gag | gag;Gag_p30\|GAG;GAG_gammaretroviridae | ERV1−3_FCa−I;LTR/ERV1 | SLC43A1;ENSFCAP00000010539 |
| Fcat62.chrD1.115549054.115549392.+ | Pro | pro;Asp\|AP;AP_pepsins_A1a | . | NA;ENSFCAP00000007246 |
| Fcat62.chrE2.7357550.7357795.+ | Pro | AP;AP_pepsins_A1a | . | NAPSA;ENSFCAP00000000296 |
| Fcat62.chrX.55076453.55076968.+ | Pro | pro;dUTPase\|DUT;DUT_caulimoviruses | (CTCTG)n;Simple_repeat | NA;ENSFCAP00000017174 |
| Fcat62.chrX.59136384.59136827.+ | Gag | gag;zf−CCHC_6 | SINEC_Fc2;SINE/tRNA | NA;ENSFCAP00000013737 |

### Pig EVE-ORFs

<table>
<thead>
<tr><th>ID</th><th>Domain</th><th>HMM</th><th>Repbase</th><th>Gene</th></tr>
</thead>
<tbody>
<tr><td>Sscr102.chr1.147292572.147293012.+</td><td>Pol</td><td>INT;INT_epsilonretroviridae|INT;INT_gammaretroviridae</td><td>KORV_I-int;LTR/ERV1</td><td>NA;ENSSSCP00000021333</td></tr>
<tr><td>Sscr102.chr1.147292864.147294852.+</td><td>Env</td><td>env;TLV_coat|ENV;ENV_deltaretroviridae|ENV;ENV_D-type_betaretroviridae|ENV;ENV_gammaretroviridae|ENV;ENV_retroviridae</td><td>KORV_I-int;LTR/ERV1</td><td>NA;ENSSSCP00000021333</td></tr>
<tr><td>Sscr102.chr2.112579816.112580115.-</td><td>Pol</td><td>INT;GIN1</td><td>.</td><td>GIN1;ENSSSCP00000026307</td></tr>
<tr><td>Sscr102.chr2.58660862.58661152.+</td><td>Pro</td><td>pro;G-patch</td><td>.</td><td>SUGP2;ENSSSCP00000022837</td></tr>
<tr><td>Sscr102.chr2.77165435.77165860.+</td><td>Pol</td><td>INT;INT_epsilonretroviridae|INT;INT_gammaretroviridae</td><td>KORV_I-int;LTR/ERV1</td><td>NA;ENSSSCP00000023068</td></tr>
<tr><td>Sscr102.chr2.77165712.77166596.+</td><td>Env</td><td>env;TLV_coat</td><td>KORV_I-int;LTR/ERV1</td><td>NA;ENSSSCP00000023068</td></tr>
<tr><td>Sscr102.chr7.132097261.132100221.-</td><td>Pro</td><td>pro;gag-asp_proteas|AP;AP_v_clade|RT;RT_17_6|RT;RT_crm|RT;RT_del|RT;RT_gypsy|RT;RT_maggy|RT;RT_pyggy|RT;RT_v_clade</td><td>(TCC)n;Simple_repeat</td><td>RTL1;ENSSSCP00000019531</td></tr>
<tr><td>Sscr102.chr7.42279971.42280228.+</td><td>Pro</td><td>AP;AP_pepsins_A1a</td><td>.</td><td>PGC;ENSSSCP00000001761</td></tr>
<tr><td>Sscr102.chr15.128593496.128594404.-</td><td>Env</td><td>env;TLV_coat</td><td>KORV_I-int;LTR/ERV1|ERV1-2_SSc-I;LTR/ERV1</td><td>NA;ENSSSCP00000023429</td></tr>
<tr><td>Sscr102.chr15.128594256.128594687.-</td><td>Pol</td><td>INT;INT_epsilonretroviridae|INT;INT_gammaretroviridae</td><td>ERV1-2_SSc-I;LTR/ERV1</td><td>NA;ENSSSCP00000023429</td></tr>
<tr><td>Sscr102.chr16.10274963.10276945.+</td><td>Pol</td><td>pol;Integrase_Zn|pol;RNase_H|pol;RVT_1|pol;RVT_thumb|INT;INT_betaretroviridae|INT;INT_deltaretroviridae|INT;INT_gammaretroviridae|RNaseH;RNaseH_betaretroviridae|RT;RT_17_6|RT;RT_alpharetroviridae|RT;RT_badnavirus|RT;RT_betaretroviridae|RT;RT_deltaretroviridae|RT;RT_gammaretroviridae|RT;RT_gmr1|RT;RT_lentiviridae|RT;RT_spumaretroviridae</td><td>ERV2-2-EC_I-int;LTR/ERVK</td><td>NA;ENSSSCP00000027525</td></tr>
<tr><td>Sscr102.chrX.95525871.95526314.+</td><td>Pol</td><td>INT;INT_epsilonretroviridae|INT;INT_gammaretroviridae</td><td>KORV_I-int;LTR/ERV1</td><td>NA;ENSSSCP00000025206</td></tr>
<tr><td>Sscr102.chrX.95526763.95528154.+</td><td>Env</td><td>env;TLV_coat|ENV;ENV_deltaretroviridae|ENV;ENV_D-type_betaretroviridae|ENV;ENV_gammaretroviridae|ENV;ENV_retroviridae</td><td>KORV_I-int;LTR/ERV1</td><td>NA;ENSSSCP00000025206</td></tr>
</tbody>
</table>

### Cow EVE-ORFs

| ID | Domain | HMM | Repbase | Gene |
|---|---|---|---|---|
| BtauUMD31.chr2.89843949.89844566.+ | LINE | pol;RVT_1 | BovB;LINE/RTE-BovB\|ART2A;SINE/RTE-BovB | AOX2;ENSBTAP00000038143 |
| BtauUMD31.chr3.54421563.54422549.+ | LINE | pol;RVT_1 | BovB;LINE/RTE-BovB\|ART2A;SINE/RTE-BovB | NA;ENSBTAP00000048119 |
| BtauUMD31.chr3.54951197.54951937.+ | LINE | pol;RVT_1 | BovB;LINE/RTE-BovB\|ART2A;SINE/RTE-BovB | NA;ENSBTAP00000019081 |
| BtauUMD31.chr4.18061526.18062050.– | LINE | pol;RVT_1 | BovB;LINE/RTE-BovB | NA;ENSBTAP00000055238 |
| BtauUMD31.chr5.29427372.29428259.+ | LINE | pol;RVT_1 | L1_BT;LINE/L1 | NA;ENSBTAP00000013453 |
| BtauUMD31.chr5.85305102.85305743.+ | LINE | pol;RVT_1 | SINE2-1_BT;SINE/tRNA\|L1_BT;LINE/L1 | CASC1;ENSBTAP00000050480 |
| BtauUMD31.chr6.14296424.14296819.– | LINE | pol;RVT_1 | BovB;LINE/RTE-BovB | ALPK1;ENSBTAP00000013457 |
| BtauUMD31.chr7.4139270.4139530.+ | Pro:Pol | pro;G-patch | . | SUGP2;ENSBTAP00000017672 |
| BtauUMD31.chr7.64723687.64725831.+ | Env | env;GP41\|ENV;ENV_B type_bertaretroviridae | ERV2-1-I_BT;LTR/ERVK\|BTLTR1F;LTR/ERVK | Fematrin-1 (BERV-K1 env) |
| BtauUMD31.chr7.104530493.104530870.– | Pol | INT;GIN1 | (AAATA)n;Simple_repeat | GIN1;ENSBTAP00000050552 |
| BtauUMD31.chr10.76487509.76487952.+ | LINE | pol;RVT_1 | ART2A;SINE/RTE-BovB\|BovB;LINE/RTE-BovB | SYNE2;ENSBTAP00000029285 |
| BtauUMD31.chr12.18173795.18174319.– | LINE | pol;RVT_1 | BovB;LINE/RTE-BovB | NA;ENSBTAP00000004630 |
| BtauUMD31.chr12.79021757.79022221.+ | LINE | pol;RVT_1 | BovB;LINE/RTE-BovB | IPO5;ENSBTAP00000043010 |
| BtauUMD31.chr13.9834956.9835924.+ | LINE | pol;RVT_1 | BovB;LINE/RTE-BovB\|ART2A;SINE/RTE-BovB | MACROD2;ENSBTAP00000045773 |
| BtauUMD31.chr13.78407524.78408978.– | Env | env;TLV_coat\|ENV;ENV_retroviridae | ERV1N-3_SSc-I;LTR/ERV1 | RUM1;ENSBTAP00000062096 |
| BtauUMD31.chr14.26441553.26442059.+ | LINE | pol;RVT_1 | BovB;LINE/RTE-BovB | SDCBP;ENSBTAP00000026526 |
| BtauUMD31.chr14.59410045.59410341.+ | LINE | pol;RVT_1 | BovB;LINE/RTE-BovB | ANGPT1;ENSBTAP00000018675 |
| BtauUMD31.chr17.54174017.54174778.– | LINE | pol;RVT_1 | ART2A;SINE/RTE-BovB\|BovB;LINE/RTE-BovB | DNAH10;ENSBTAP00000029662 |
| BtauUMD31.chr18.29556085.29558388.– | LINE | pol;RVT_1 | L1_BT;LINE/L1 | CDH8;ENSBTAP00000030891 |
| BtauUMD31.chr19.40030158.40030817.– | LINE | pol;RVT_1 | ART2A;SINE/RTE-BovB\|BovB;LINE/RTE-BovB | CWC25;ENSBTAP00000040794 |
| BtauUMD31.chr21.55339309.55339986.+ | LINE | pol;RVT_1 | BovB;LINE/RTE-BovB\|ART2A;SINE/RTE-BovB | NA;ENSBTAP00000040927 |
| BtauUMD31.chr21.67427517.67431587.– | Pro:Pol | pro;gag-asp_proteas\|GAG;GAG_v_clade\|RT;RT_17_6\|RT;RT_crm\|RT;RT_del\|RT;RT_gypsy\|RT;RT_maggy\| RT;RT_pyggy\|RT;RT_v_clade | . | RTL1;ENSBTAP00000055386 |
| BtauUMD31.chr22.11528351.11529376.+ | LINE | pol;RVT_1 | BovB;LINE/RTE-BovB\|BTLTR1;LTR/ERVK | DLEC1;ENSBTAP00000011298 |
| BtauUMD31.chr22.55015756.55017120.– | LINE | pol;RVT_1 | ART2A;SINE/RTE-BovB\|BovB;LINE/RTE-BovB\|BTLTR1;LTR/ERVK | ATP2B2;ENSBTAP00000001723 |
| BtauUMD31.chr24.12817236.12819377.+ | Env | env;GP41\|ENV;ENV_B type_bertaretroviridae | ERV2-1-I_BT;LTR/ERVK\|ERV2-1-LTR_BT;LTR/ERVK | BERV-K2 env |
| BtauUMD31.chr25.19468570.19469076.+ | LINE | pol;RVT_1 | BovB;LINE/RTE-BovB | NA;ENSBTAP00000008832 |
| BtauUMD31.chr28.12921485.12921745.+ | LINE | pol;RVT_1 | BovB;LINE/RTE-BovB | ZNF33B;ENSBTAP00000024349 |
| BtauUMD31.chrX.84544746.84545081.– | Gag | gag;zf-CCHC_6 | . | NA;ENSBTAP00000053806 |

### Platypus EVE-ORFs

| ID | Domain (HMM) | Repbase | Gene |
|---|---|---|---|
| Oana5.chrX1.32227928.32228290.+ | pol;RVT_1 | Mon1d;SINE/MIR\|L2_Plat1m;LINE/L2 | FGD5;ENSOANP00000017248 |

**Table S2** Species and Genome assembly used in this study.

Genome ID: Species name in the gEVE database (http://geve.med.u-tokai.ac.jp)

| Species | Nomenclature | Genome Database | Genome ID |
|---|---|---|---|
| Human | Homo sapiens | GRCh38, Dec 2013 | Hsap38 |
| Chimpanzee | Pan troglodytes | CSAC 2.1.4/panTro4, Feb 2011 | Ptro214 |
| Gorilla | Gorilla gorilla gorilla | gorGor3.1/gorGo3, May 2011 | Ggor31 |
| Orangutan | Pongo pygmaeus abelii | PPYG2, Sep 2007 | Pabe2 |
| Baboon | Papio anubis | Panu_2.0, Jun 2012 | Panu2 |
| Macaque | Macaca mulatta | MMUL 1.0, Feb 2006 | Mmul1 |
| Marmoset | Callithrix jacchus | C_jacchus3.2.1, Jan 2010 | Cjac321 |
| Mouse | Mus musculus | GRCm38.p1, Jan 2012 | Mmus38 |
| Rat | Rattus norvegicus | Rnor_5.0, Mar 2012 | Rnor50 |
| Rabbit | Oryctolagus cuniculus | oryCun2, Nov 2009 | Ocun2 |
| Cow | Bos taurus UMD3.1 | UMD3.1, Dec 2009 | BtauUMD31 |
| Cow | Bos taurus 4.6.1 | Btau_4.6.1 Nov 2011 | Btau461 |
| Dog | Canis lupus familiaris | CanFam3.1, Sep 2011 | Cfam31 |
| Cat | Felis catus | Felis_catus_6.2, Sep 2011 | Fcat62 |
| Horse | Equus caballus | Equ Cab 2, Sep 2007 | Ecab2 |
| Sheep | Ovis aries | Oar_v3.1, 2012/09/24 | Oari31 |
| Pig | Sus scrofa | Sscrofa10.2, Aug 2011 | Sscr102 |
| Goat | Capra hircus | CHIR_1.0, Jan 2013 | Chir1 |
| Opossum | Monodelphis domestica | monDom5, Oct 2006 | Mdom5 |
| Platypus | Ornithorhynchus anatinus | OANA5, Dec 2005 | Oana5 |

**Table S3** Accession numbers of runs used in myoblast differentiation analysis.

**A)** Human, **B)** Mouse. NOTE: We did not use a mouse replicate SRA data for day3 because it was broken.

**A** Human (SRP033135)

| Run ID | BioSample | Sample name | Experiment | Instrument | Type in this stuc | Library protocol | Label |
|---|---|---|---|---|---|---|---|
| SRR1033282 | SAMN02413273 | GSM1269332 | SRX379972 | Illumina HiSeq 2500 | D0 | Bulk RNA-Seq (TruSeq) | Bulk RNA-Seq_0 hrs |
| SRR1033283 | SAMN02413261 | GSM1269333 | SRX379973 | Illumina HiSeq 2500 | D0 | Bulk RNA-Seq (TruSeq) | Bulk RNA-Seq_0 hrs |
| SRR1033284 | SAMN02413275 | GSM1269334 | SRX379974 | Illumina HiSeq 2500 | D0 | Bulk RNA-Seq (TruSeq) | Bulk RNA-Seq_0 hrs |
| SRR1033285 | SAMN02413262 | GSM1269335 | SRX379975 | Illumina HiSeq 2500 | D1 | Bulk RNA-Seq (TruSeq) | Bulk RNA-Seq_24 hrs |
| SRR1033286 | SAMN02413276 | GSM1269336 | SRX379976 | Illumina HiSeq 2500 | D1 | Bulk RNA-Seq (TruSeq) | Bulk RNA-Seq_24 hrs |
| SRR1033287 | SAMN02413264 | GSM1269337 | SRX379977 | Illumina HiSeq 2500 | D1 | Bulk RNA-Seq (TruSeq) | Bulk RNA-Seq_24 hrs |
| SRR1033288 | SAMN02413277 | GSM1269338 | SRX379978 | Illumina HiSeq 2500 | D2 | Bulk RNA-Seq (TruSeq) | Bulk RNA-Seq_48 hrs |
| SRR1033289 | SAMN02413263 | GSM1269339 | SRX379979 | Illumina HiSeq 2500 | D2 | Bulk RNA-Seq (TruSeq) | Bulk RNA-Seq_48 hrs |
| SRR1033290 | SAMN02413278 | GSM1269340 | SRX379980 | Illumina HiSeq 2500 | D2 | Bulk RNA-Seq (TruSeq) | Bulk RNA-Seq_48 hrs |
| SRR1033291 | SAMN02413265 | GSM1269341 | SRX379981 | Illumina HiSeq 2500 | D3 | Bulk RNA-Seq (TruSeq) | Bulk RNA-Seq_72 hrs |
| SRR1033292 | SAMN02413279 | GSM1269342 | SRX379982 | Illumina HiSeq 2500 | D3 | Bulk RNA-Seq (TruSeq) | Bulk RNA-Seq_72 hrs |
| SRR1033293 | SAMN02413266 | GSM1269343 | SRX379983 | Illumina HiSeq 2500 | D3 | Bulk RNA-Seq (TruSeq) | Bulk RNA-Seq_72 hrs |

**B** Mouse (SRP036149)

| Run ID | BioSample | Sample name | Experiment | Instrument | Type in this stuc | LibrarySelection | Label |
|---|---|---|---|---|---|---|---|
| SRR1156540 | SAMN02615744 | day0_rep1 | SRX460743 | Illumina HiSeq 2500 | D0 | PolyA | day0_rep1 |
| SRR1156548 | SAMN02615744 | day0_rep1 | SRX460743 | Illumina HiSeq 2500 | D0 | PolyA | day0_rep1 |
| SRR1156614 | SAMN02615745 | day0_rep2 | SRX460803 | Illumina HiSeq 2500 | D0 | PolyA | day0_rep2 |
| SRR1156621 | SAMN02615745 | day0_rep2 | SRX460803 | Illumina HiSeq 2500 | D0 | PolyA | day0_rep2 |
| SRR1156654 | SAMN02615746 | day0_rep3 | SRX460855 | Illumina HiSeq 2500 | D0 | PolyA | day0_rep3 |
| SRR1156816 | SAMN02615746 | day0_rep3 | SRX460855 | Illumina HiSeq 2500 | D0 | PolyA | day0_rep3 |
| SRR1156931 | SAMN02615747 | day3_rep1 | SRX461126 | Illumina HiSeq 2500 | D3 | PolyA | day3_rep1 |
| SRR1156937 | SAMN02615747 | day3_rep1 | SRX461126 | Illumina HiSeq 2500 | D3 | PolyA | day3_rep1 |
| SRR1156940 | SAMN02615748 | day3_rep2 | SRX461143 | Illumina HiSeq 2500 | D3 | PolyA | day3_rep2 |
| SRR1156941 | SAMN02615748 | day3_rep2 | SRX461143 | Illumina HiSeq 2500 | D3 | PolyA | day3_rep2 |
| SRR1156943 | SAMN02615749 | day3_rep3 | SRX461144 | Illumina HiSeq 2500 | D3 | PolyA | day3_rep3 |
| SRR1156944 | SAMN02615750 | day6_rep1 | SRX461145 | Illumina HiSeq 2500 | D6 | PolyA | day6_rep1 |
| SRR1156945 | SAMN02615750 | day6_rep1 | SRX461145 | Illumina HiSeq 2500 | D6 | PolyA | day6_rep1 |
| SRR1156946 | SAMN02615751 | day6_rep2 | SRX461146 | Illumina HiSeq 2500 | D6 | PolyA | day6_rep2 |
| SRR1156947 | SAMN02615751 | day6_rep2 | SRX461146 | Illumina HiSeq 2500 | D6 | PolyA | day6_rep2 |
| SRR1156948 | SAMN02615752 | day6_rep3 | SRX461147 | Illumina HiSeq 2500 | D6 | PolyA | day6_rep3 |
| SRR1156949 | SAMN02615752 | day6_rep3 | SRX461147 | Illumina HiSeq 2500 | D6 | PolyA | day6_rep3 |
